# SARS-CoV-2 membrane protein conformations induce distinct membrane curvatures

**DOI:** 10.64898/2026.07.06.736641

**Authors:** Joseph McTiernan, Roya Zandi, Michael E. Colvin, Ajay Gopinathan

## Abstract

The assembly and budding of enveloped viruses requires thousands of membrane proteins to collectively remodel host-cell membranes into highly curved virions. In SARS-CoV-2, this process is driven by interactions between viral structural proteins and the endoplasmic reticulum–Golgi intermediate compartment (ERGIC) membrane. The membrane (M) protein, an embedded homodimer and the most abundant viral component, exists in two conformations: a compact “short” form and an elongated “long” form. Although M is essential for virion assembly, how its conformations contribute to the generation and organization of the membrane curvature required for budding has remained unknown. Here, we used all-atom and Martini coarse-grained molecular dynamics simulations to show that individual M proteins can induce distinct membrane curvatures, depending on their conformation. The long form bends the membrane around its C-terminal, forming a valley-like depression, while the short form predominantly bends the membrane away from the C-terminal producing an anisotropic ridge. The induced curvatures correspond to the bulb and neck regions of a budding virion, respectively. Coarse-grained simulations of M protein pairs further reveal that curvature modulates long-range, membrane-mediated M–M interactions, leading to repulsion between dissimilar conformations. Together, these results suggest that the long and short forms of M naturally segregate to shape the virion’s bulb and neck, potentially facilitating genome encapsulation and membrane scission. This mechanism provides a physical basis for coronavirus budding and suggests that conformationally encoded curvature fields may represent a general principle underlying the formation of enveloped viruses.

**Significance Statement:** Enveloped viruses must bend host cell membranes to form new viral particles, but the driving force for membrane bending has been unclear. Using molecular dynamics simulations, we show that the SARS-CoV-2 membrane protein, the most abundant structural protein of the virus, drives membrane bending, and that its two natural conformations generate opposite signs of curvature which match the geometry of distinct regions of a budding virus. Moreover, these induced curvatures cause proteins in different conformations to repel one another providing a physical basis for their spatial segregation. Our results suggest that the membrane protein’s conformations can contribute significantly to the membrane remodeling needed for virus assembly and release, revealing a simple mechanism that may apply to other enveloped viruses.

---

**T**he remodeling of biological membranes into highly curved structures is fundamental to processes ranging from intracellular trafficking and organelle biogenesis to cell division. This problem is especially striking during the assembly of enveloped viruses, where thousands of viral proteins must coordinate membrane remodeling to generate a highly ordered virion. Coronavirus serves as a compelling instance of this process. Its virions consist of a positive single-stranded RNA genome enveloped by the host cell’s membrane, encompassing four main structural proteins: the membrane (M) protein, nucleocapsid (N) protein, envelope (E) protein, and spike (S) protein (1). Each viral protein plays a vital role in replication, with the S protein responsible for binding and entry into the host cell (2, 3). Following entry and translation, the M, E, and S proteins localize to the endoplasmic reticulum-to-Golgi intermediate compartment (ERGIC) membrane, while N protein and viral RNA accumulate within the adjacent cytoplasm (4, 5). The most abundant structural protein is the M protein which guides E and S proteins along the membrane, leading to the formation of curvature-inducing protein clusters (6–8). Simultaneously, the N protein recruits viral RNA (vRNA), forming a complex that then binds to M protein along the membrane (7, 9–11). Lastly, the E protein is a viroporin that helps retain S protein (12), and may induce membrane curvature or close the neck of the virion to complete virion formation (13–15). The interactions of these structural proteins with each other and the membrane drives viral assembly and budding, resulting in virion formation.

These functions are largely conserved across coronaviruses, with the M protein always required for complete assembly and budding (12, 16–18). This essential role presumably arises from its ability to form protein clusters with other viral components across the membrane and bind to N/vRNA complexes during assembly and budding (6–11) as well as its capacity for higher order oligomerization (7, 19, 20). In general, the membrane curvature necessary for budding can be generated by the wrapping of compact genomic complexes and/or the induction of curvature by membrane embedded protein complexes. In fact, transmembrane proteins commonly induce curvature in membranes driven by their shape asymmetry (21, 22). Membrane-mediated interactions between proteins due to induced curvature have also been characterized through experimental (23, 24), analytical (25–28), and computational (23, 29, 30) studies. Although M protein, with its roughly conical shape, is a good candidate for generating curvature required for virion assembly and budding, the molecular origins of curvature generation in coronaviruses remain unclear.

Recent cryo-electron microscopy studies have determined the Severe Acute Respiratory Syndrome Coronavirus 2 (SARS-CoV-2) M protein is a homodimer with two conformations, an elongated or ‘long’ form (19) and a compact ‘short’ form (19, 31). Each conformation consists of an intravirion C-terminal domain, a transmembrane domain with three helices, and an extravirion N-terminal domain (19). Observations of the SARS-CoV virion showed that the longform interacts with the N/RNA complex (via the C-terminal) and preferentially occupied regions of curvature consistent with the virion bulb while the short form preferred thinner membrane regions (7). More recently it has been shown that the existence of both conformations throughout assembly and budding is vital, with virion formation inhibited by a small molecule that stabilized the SARS-CoV-2 M protein in its short form (20). This was attributed to the supposed insufficient production of membrane curvature by the short form and the reduced oligomerization caused by the molecule. Similarly, another molecule capable of transitioning the M protein to an unnatural conformation also inhibited assembly (32). Studies investigating the effects of the SARS-CoV-2 M protein on the membrane showed that the short form thinned the surrounding membrane (33, 34), leading to a line tension acting as a weak membrane-mediated protein-protein attraction. The membrane deformations reported there were suggestive of curvature generation but were not conclusive, and no further quantification of induced curvature for either conformation has been performed.

To determine the existence of conformation dependent curvature and its effect on membrane-mediated interactions, we utilize all-atom and Martini 3 coarse-grained (CG) molecular dynamics (MD) simulations of SARS-CoV-2 M proteins embedded in an ERGIC-mimetic lipid bilayer membrane. All-atom simulations allow for a more refined representation of protein induced membrane curvature (35–39) while coarse-graining the system permits simulation of larger time scales, multiple proteins and membrane-mediated interactions (37, 38, 40–45). Using these simulations, we show that individual M protein conformations in ERGIC-mimetic membranes generate distinct curvatures, with the long form bending the membrane toward its C-terminal consistent with the virion bulb curvature and the short form bending the membrane away from its C-terminal along one axis to produce a ridge-like curvature corresponding to the virion neck curvature. This is consistent with the substantial kink that develops in the long form’s transmembrane domain, reversing the orientation of its conical shape relative to the short form. Extending our simulations to two-protein systems, we find that curvature drives long-range membrane-mediated interactions, with like conformations showing a tendency to attract while unlike conformations tend to repel. Taken together, our results show that M protein curvature generation both shapes the budding membrane and drives membrane-mediated interactions that segregate conformations into geometrically defined regions of the bud. Our results are consistent with *in vitro* observations (7) of different membrane curvature preferences for different M protein conformations and in agreement with experiments (20) showing that the short form alone was insufficient for assembly and budding, suggesting the crucial role conformational polymorphism in a single viral protein plays.

## Results

### Curvature induction of the SARS-CoV-2 membrane protein is conformation dependent

The assembly and budding process for SARS-CoV-2 involves the M protein acting as a membrane embedded matrix protein, guiding the formation of potentially curvature inducing protein clusters to drive budding, as shown in *SI Appendix*, Fig. S1. To determine whether M protein can be responsible for inducing the curvature required for viral assembly and budding, we performed all-atom and Martini CG MD simulations of single SARS-CoV-2 M protein conformations embedded in an ERGIC-like 40 nm x 40 nm lipid bilayer (Fig. 1A). To measure the deformation induced by the protein, we quantify the average membrane midplane height, corresponding to the mean displacement (from reference values along the membrane boundary) of the upper and lower leaflets, averaged over time. Fig. 1B is a schematic showing a protein inducing a negative deformation (negative membrane midplane height) with the membrane bending toward the C-terminal. The surface curvature, related to the second derivative of the membrane height, is positive in this case.

**Fig. 1.**
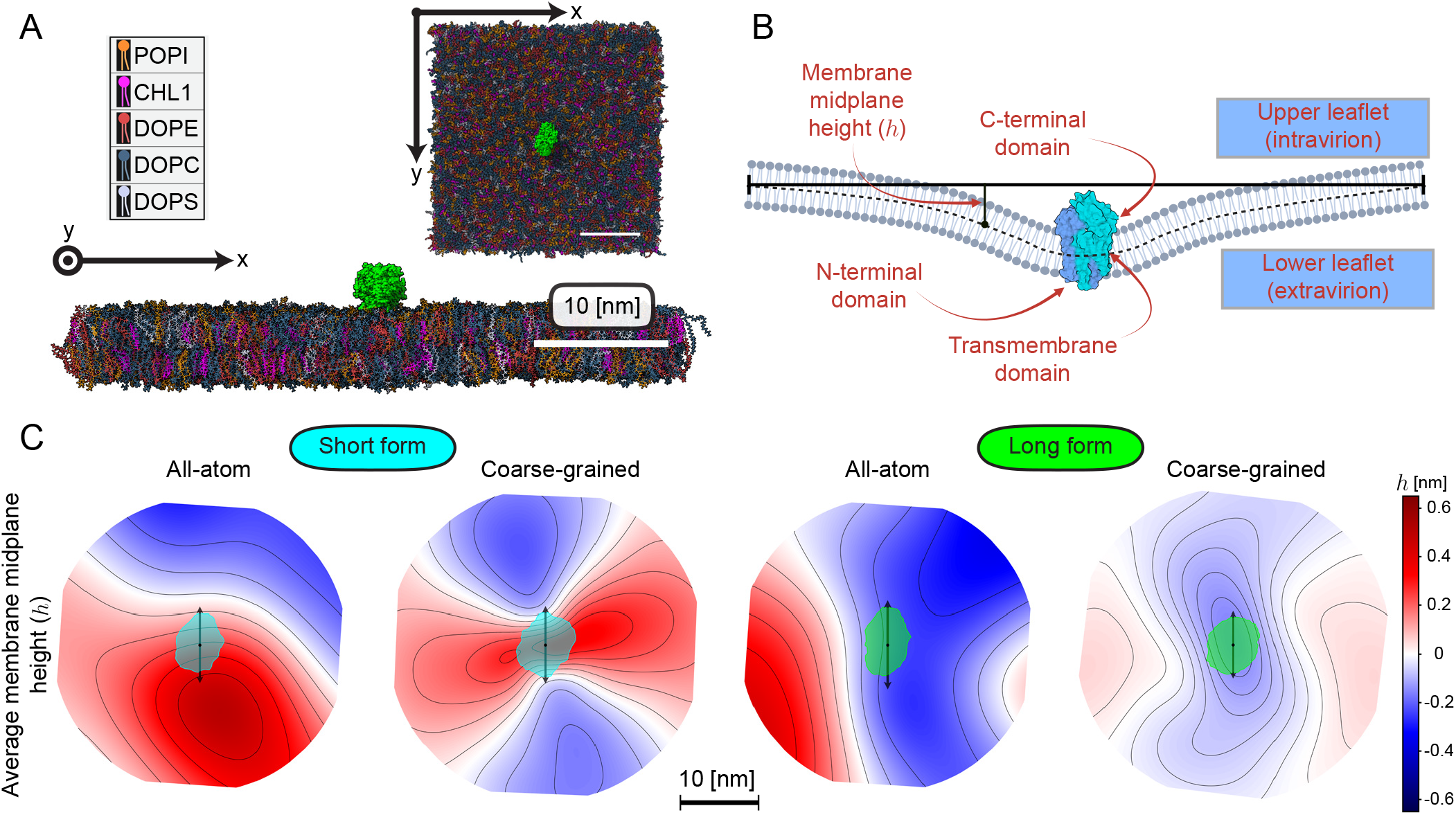
Average membrane midplane height from all-atom and coarse-grained MD simulations show that induced deformations are conformation dependent. (A) View from above and the side of an initial all-atom simulation frame with the long form (green) embedded in a 40 nm x 40 nm lipid bilayer physiologically similar to the ERGIC. The lipid bilayer is colored according to composition, where the scale-bar corresponds to 10 nm, and the protein’s C-terminal is on top. (B) Cartoon characterizing the M protein’s embedding, where the membrane midplane height is defined relative to the average value along the edge of the membrane and the C-terminal domain is inside the virion. (C) Average membrane midplane height in the protein’s reference frame for each conformation, where we averaged from 250 ns to 750 ns for all-atom and 2 *µ*s to 30 *µ*s for Martini CG simulations. Each mesh is scaled identically. The protein’s cumulative cross-section (over simulation time) is shown in cyan for the short form and green for the long form. Black dots indicate the protein’s average center of mass while black double-sided arrows show the orientation of its major axis in the plane of the membrane. Meshes are oriented such that these axes are vertical for visualization. BioRender.com was used to create the legend for (A) and the schematic in (B).

To quantify induced deformations, we used radial basis interpolation from raw simulation data to fit polynomials describing the average membrane midplane height in the x-y plane with respect to the protein’s reference frame (Fig. 1C). This procedure provides a continuous surface along the simulation box, allowing for more accurate deformation and curvature calculations relative to the protein’s orientation in the membrane. It should be noted that not accounting for protein rotation during trajectory processing leads to a radially smeared height that is continuous across simulation boundaries, as shown in *SI Appendix*, Fig. S2. Examining these surfaces in the protein’s reference frame, we see that the membrane deformation changes sign with conformation. The short form induces a positive deformation with the membrane bending away from the C-terminal, while the long form induces a negative deformation with the membrane bending toward the C-terminal. While all-atom and CG simulations give similar sign, the scale differs slightly, with the short form maximum deformation changing from ∼0.50 nm in all-atom simulations to ∼0.34 nm in our CG counterpart. Similarly, the long form’s induced maximum deformation is on the scale of ∼ −0.36 nm and ∼ −0.16 nm for all-atom and CG simulations respectively. This scale deviation likely reflects several factors, including a shorter all-atom sampling time, and differences between atomistic and coarse-grained membrane elasticity properties. We note that Gaussian smoothing with a standard deviation comparable to bilayer thickness for all-atom height was used to somewhat limit this effect (*SI Appendix*, Sec. 1, results without smoothing can be seen in *SI Appendix*, Fig. S3). Given this shorter sampling time, we predominantly use all-atom simulations to corroborate the trends observed in our CG systems as opposed to providing higher spatial resolution characterizations.

Finally, to directly verify that the observed deformations are induced by the protein as opposed to the protein localizing in regions with membrane fluctuations of preferred curvature, we conducted 10 *µ*s long CG simulations of individual short and long conformations with each restrained to maintain their initial x-y center of mass. These results were consistent with the earlier simulations (*SI Appendix*, Fig. S4), confirming that the deformations induced by the short and long forms are positive and negative respectively.

### The short form induces a sharp ridge with the C-terminal on top, while the long form induces a smoother valley

Next, we determined the magnitude and spatial profile of the induced curvature of a single M protein relative to its orientation in the plane of the membrane. Using the CG (2 *µs* to 30 *µs*) average membrane midplane polynomials, with corresponding meshes shown in Fig. 1C, we obtained the maximum (Fig. 2A) and minimum curvatures (Fig. 2B) at each point along the surface using the shape operator. This operator allows for quantification of both magnitude (*k*_1_, *k*_2_) and direction 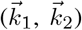 of the maximum and minimum curvatures at a point on the surface, which are defined as the largest and smallest curvatures of all curves passing through the point. The protein’s induced, or spontaneous, maximum and minimum curvatures shown were obtained by averaging values adjacent to its center of mass.

**Fig. 2.**
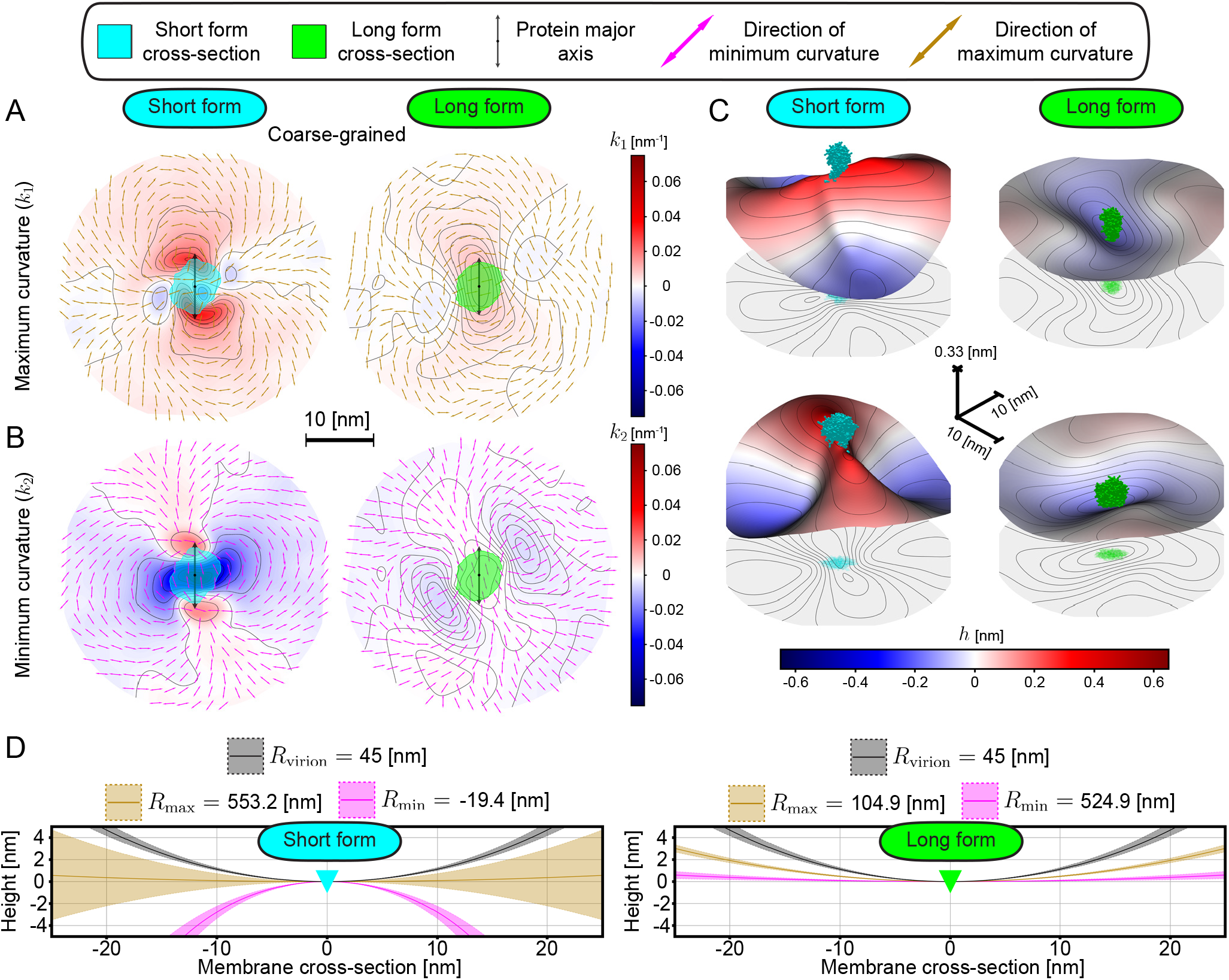
SARS-CoV-2 M protein induced membrane curvature fields from MD simulations show that the short form induces a ridge, while the long form induces a valley. (A) Maximum (*k*_1_ ) and (B) minimum (*k*_2_ ) curvatures are shown in the reference frame of each conformation for coarse-grained simulations. A positive curvature represents a negative deformation, or the membrane curling around the C-terminal. The protein’s cumulative cross-section is colored cyan for the short form and green for the long form, where its major axis is oriented vertically as a black double-sided arrow. Isolines for the corresponding scalar quantities are black lines along the surface. For (A) and (B), the local direction of maximum and minimum curvature is represented with brown and pink double-sided arrows respectively. At each point, 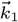 and 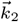 are orthogonal, with *k*_1_ > *k*_2_ . (C) Different three-dimensional visualizations of average membrane midplane height are displayed for coarse-grained simulations of both conformations, where the height of the mesh is scaled by a factor of 30. Each conformation’s initial, unscaled solvent excluded surface is embedded and colored cyan for the short form and green for the long form. The projection of this resulting mesh is embedded below. (D) Visual representation of membrane cross-sectional profiles as lines with different radii of curvature corresponding to minimum and maximum curvatures generated by the short and long forms. Proteins are represented as triangles with the C-terminal as the base. Shading represents error. Black curves correspond to the virion’s radius of curvature (*R*_virion_ = 45 nm ± 5 nm), while brown and pink correspond to the radius of maximum and minimum curvature respectively.

The induced maximum and minimum curvatures of the short form are significantly different, with the maximum curvature approximately zero (0.002 ± 0.013 nm^−1^) and the minimum curvature a much larger negative quantity ( −0.052 ±0.009 nm^−1^). Reported uncertainties account for both methodology driven intrinsic error and spatial variations surrounding the protein’s center of mass. Fig. 2A and Fig. 2B suggest that the short form generates a ridge, where its major axis lies roughly perpendicular to the ridge length, and parallel to the ridge width. The long form’s induced maximum curvature is 0.010± 0.001 nm^−1^ and the minimum curvature is 0.002 ± 0.001 nm^−1^. Thus, the induced curvatures of the long form are positive in both principal directions and not as anisotropic as that of the short form’s, suggesting that the long form induces a valley. Radial averages of the curvature direction adjacent to the protein’s center of mass allow us to quantify the protein’s orientation relative to the membrane deformation, showing that each protein’s major axis appears to coincide with the direction of minimum curvature (*SI Appendix*, Fig. S5). An anisotropic curvature like that induced by the short form coupled to the protein’s orientation could help define protein-protein interactions, influencing alignment between them.

Three-dimensional visualizations of the induced surface for both M protein conformations, displayed in Fig. 2C, provide two different views, one pointing along the y-axis (top), and the other along the x-axis (bottom). These representations confirm that the long form induces a valley while the short form induces a ridge. In Fig. 2D, we include a visual representation of the magnitude and sign of each conformation’s induced principal curvatures and compare them to the virion radius of curvature. We note that the short form induced curvature is very sharp with a radius less than half the virion radius and of the opposite sign. On the other hand, while the long form curvature sign is consistent with that of the virion, the long form’s smallest radius of curvature is still approximately a factor of two larger than the virion radius.

### The long form curvature profile is consistent with a budding virion, whereas the short form’s matches the neck

Total curvature (2*C* = *k*_1_ + *k*_2_) serves as a metric for curvature generation independent of protein orientation in the plane of the membrane. Note that 2*C* > 0 indicates that the membrane tends to curl around the protein’s C-terminal, while 2*C* < 0 implies a preference to curl away. To estimate the total curvature that a protein induces, we calculated the average value of *k*_1_ + *k*_2_ in the region around the protein’s center of mass, and plot it for both short and long forms under both free and restrained conditions in Fig. 3A. Restraining the protein did not qualitatively alter the resulting membrane shape (*SI Appendix*, Fig. S4), with short and long forms continuing to induce negative and positive total curvature respectively. Similarly, the scale and direction of induced minimum and maximum curvatures remained comparable between restrained and free short forms (*SI Appendix*, Fig. S5 and S6). However, for the restrained long form, the direction of minimum curvature was shifted slightly away from its major axis with an enhanced induced maximum curvature (*SI Appendix*, Fig. S5 and S6).

**Fig. 3.**
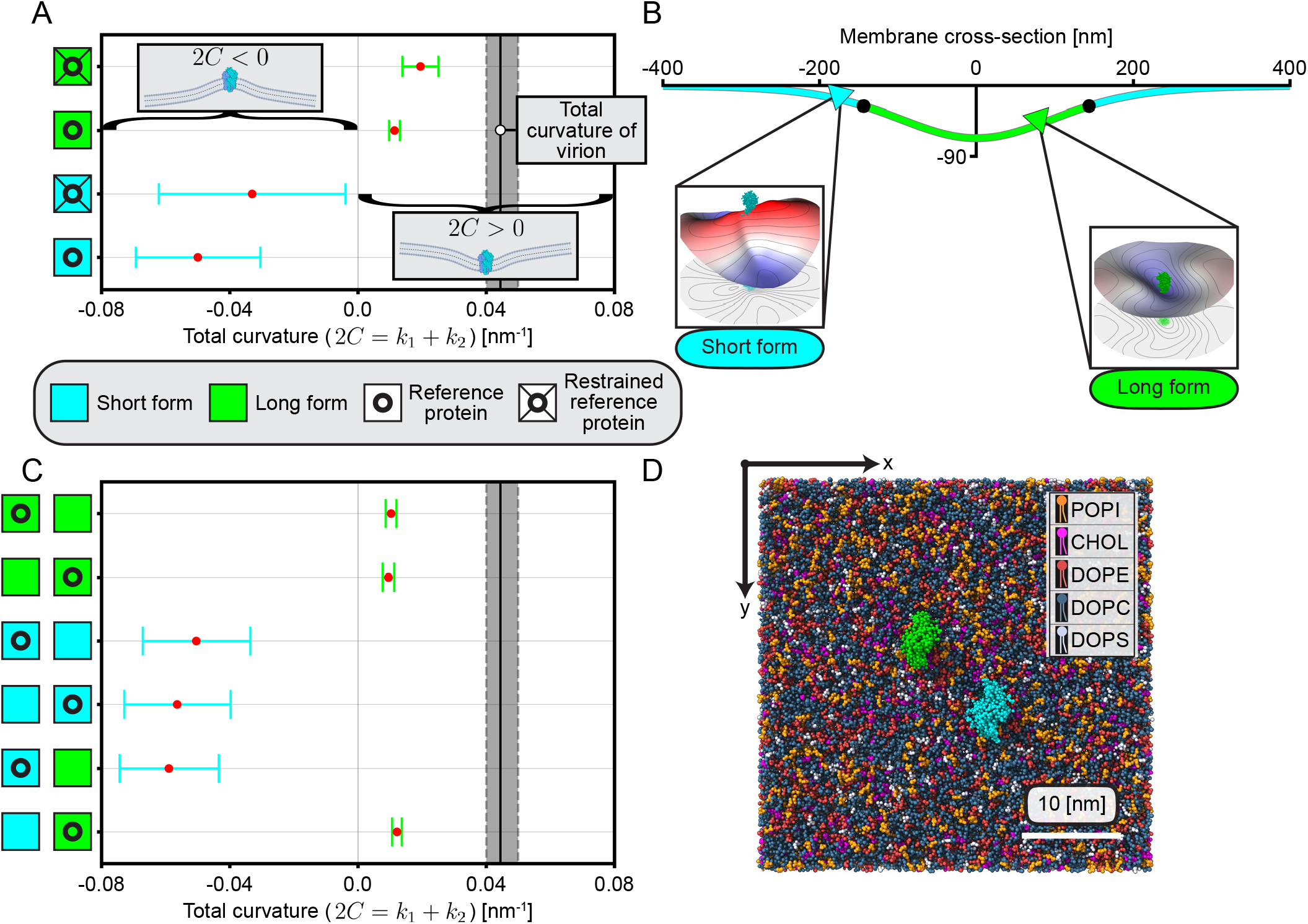
The long form curvature profile matches the center of a budding virion while the short form is consistent with the neck, with neither influenced by introducing another conformation. (A) Total curvature (2*C* = *k*_1_ + *k*_2_ ) calculated from each single conformation CG simulation, where cyan and green squares represent the short and long forms respectively. Local average total curvature of the reference protein is plotted with a red dot. Simulations with free proteins are labeled with a square surrounding a circle while restrained protein (in the x-y plane) simulations are identified with lines connecting the circle to each corner of the square. Error for every quantity is displayed as total curvature spread. The solid, vertical black line defines the virion total curvature corresponding to *R*_virion_ = 45 nm ± 5 nm. (B) Membrane cross-section of a budding virion, with cyan (green) corresponding to regions with negative (positive) total curvature and black dots indicating zero curvature. The short and long forms are represented with cyan and green triangles respectively. Each triangle’s base corresponds to the conformation’s C-terminal. Insets: three-dimensional visualizations of proteins and surrounding membrane shape. (C) Total curvature in the region around the reference protein is calculated for each two-protein CG simulation and shown as a red dot. (D) CG simulation schematic for beginning of typical two-protein simulation, with one long form (green) and one short form (cyan) embedded in the membrane. Protein centers of mass are separated by 10 nm along the box diagonal, and principal axes are parallel. BioRender.com was used to create the insets in (A) and the legend for (D).

Comparison to the vertical black line corresponding to a fully formed virion’s total curvature in Fig. 3A suggests that neither a single short nor long form M protein generates sufficient curvature to be solely responsible for the final virion shape. In particular, while the long form induced maximum curvature is within a factor of two of the virion’s curvature (Fig. 2D), the total curvature difference is more pronounced due to the induced minimum curvature being a small positive value. Furthermore, curvature generated by a single M protein is local, and should not be viewed as sufficient to produce the final curvature of a mature virion. Full virion-scale curvature likely requires collective effects from multiple M proteins and additional assembly components. Nevertheless, the curvatures individually induced by these conformations are comparable to those that may occur during the initial stages or onset of budding.

To highlight this point, we first consider a potential membrane cross-section shape using an ansatz from (46) with parameters tuned to obtain a shape consistent with the onset of budding (see Fig. 3B). We note that this shape is not an exact solution to the membrane shape equation, and is solely being used for illustrative purposes. The curvatures of the two forms (from Fig. 3A) suggest that the long form would preferentially localize in the head of the forming bud, whereas the short form would do so along the edge or neck as shown in Fig. 3B. We note that repeating this characterization using each protein’s induced Gaussian curvature (*K* = *k*_1_*k*_2_), leads to similar localization preferences for both conformations (*SI Appendix*, Fig. S7). This localization pattern further suggests that the long form has a more central role in forming the bud while the short form promotes the formation of the neck leading to eventual scission.

### Curvature induction governs membrane-mediated M-M interactions

Next, to identify the impact of curvature induction on M-M interactions, we performed CG simulations involving the three possible conformation combinations (short-short, long-long, and long-short). Independent of the system, measuring curvatures within a protein’s reference frame shows both short and long forms consistently induce the expected negative and positive total curvature even in the presence of another protein (Fig. 3C-D). Furthermore, each measured minimum and maximum curvature - direction included - remains consistent with our individual protein results (*SI Appendix*, Fig. S5, S6, and S8). It is possible that a higher protein density is required to see a collective change in curvature induction, as described in (25).

By classifying the motion of these proteins according to their separation and relative alignment (defined in Fig. 4A), we aimed to identify the effect of curvature on M-M interactions. For the simulation with two short forms, we see initial oscillatory behavior in separation (Fig. 4B) up to approximately 13 *µs*, ranging from ∼5 nm to ∼25 nm. The proteins then bind to each other and remain so for ∼3 *µs*, where they then return back to oscillation - while also achieving another ∼1 *µs* more sporadic binding event at the ∼20 *µs* mark. In contrast, the simulation with a short and long form shows the conformations remaining largely separate, with the proteins typically ∼20 nm apart outside of a ∼1 *µs* long contact event. This is one indicator of an effective repulsion driven by induced curvature due to differing curvature profiles. The case with two long forms is more balanced with the separation between the proteins alternating between ∼10 nm and ∼20 nm, and no significant contact event is observed. Overall we see that like conformations remain closer than dissimilar ones, which repel, with contact events only occurring for simulations with a short form. Furthermore, binding events tend to coincide with proteins being anti-aligned (Fig. 4B and *SI Appendix*, Fig. S9). One dimensional histograms for both separation and alignment in these simulations further emphasize these conclusions (*SI Appendix*, Fig. S10), where anti-alignment between two short forms arises due to the pronounced binding event. Note that interactions with periodic images become prominent at large separations and prevent further separation.

**Fig. 4.**
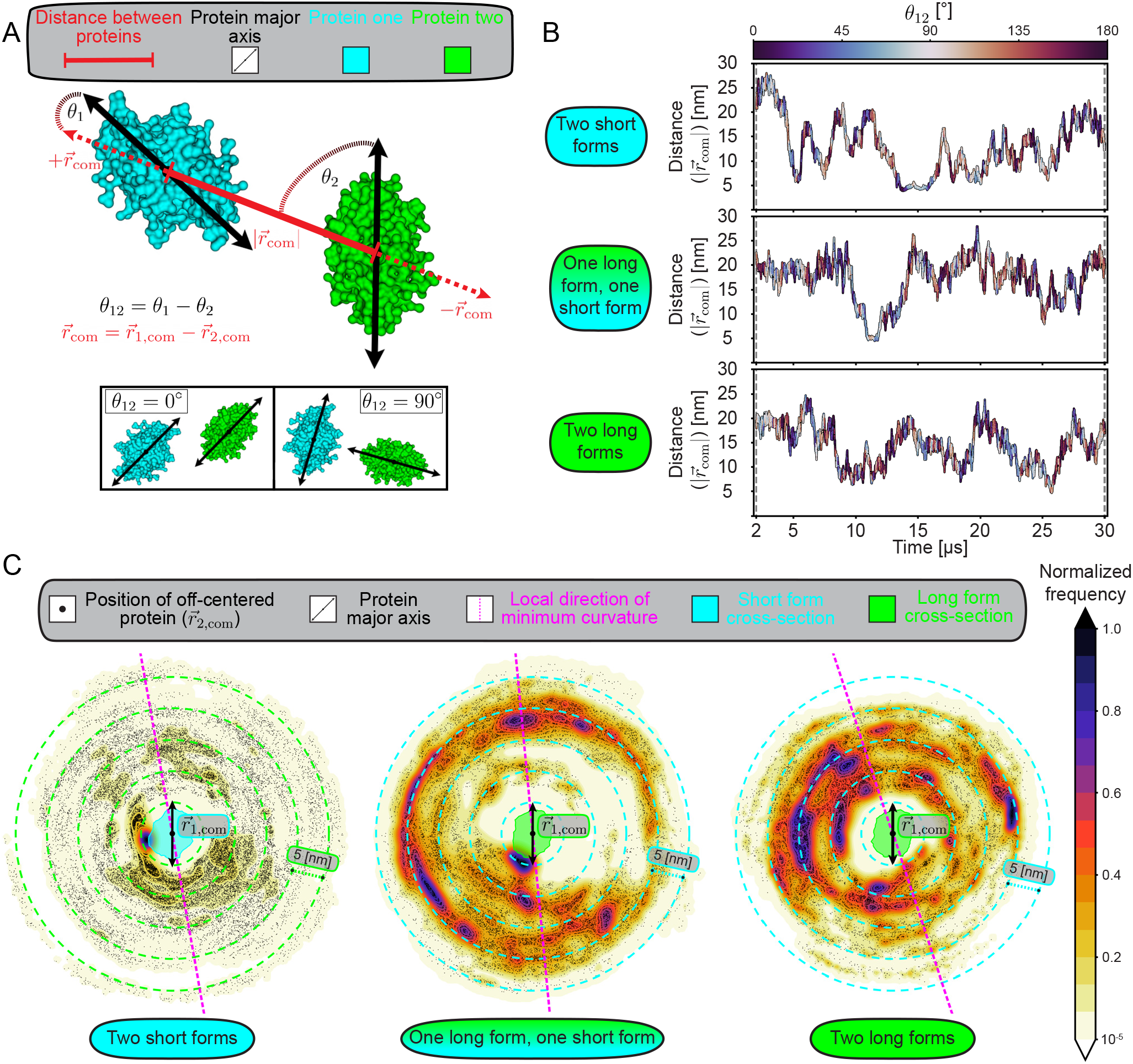
Curvature induction is capable of separating and aligning M proteins. (A) Schematic defining the separation and alignment between two proteins, with the solvent excluded surface area of protein one colored cyan and that of protein two colored green. Protein one denotes the reference protein. Distance, or separation, between proteins is defined according to the center of mass vector between them (red, 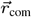 ). The angle each protein’s major axis (black double-sided arrow) makes with 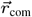is *θ*_i_, where alignment between proteins is *θ*_12_ = *θ*_1_ −*θ*_2_ . All angles are modulo 180^*°*^, with alignment *θ*_12_ = 0^*°*^ and anti-alignment *θ*_12_ = 90^*°*^. (B) Separation between proteins in each multiple protein simulation, with curves starting at 2 *µs*, ending at 30 *µs*, and colored according to alignment. (C) Localization of protein two in protein one’s reference frame, colored according to its frequency normalized with respect to the most common position. Center of mass for the off-centered protein at each point in time is represented with a black dot, while the reference protein’s cumulative cross-section is shown in cyan (short form) or green (long form). Each plot is fixed so the major axis of the reference protein points vertically, indicated by the double headed arrow with protein center of mass in the middle. Pink dotted lines correspond to local 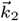, while cyan or green dotted lines indicate radial distances in units of 5 nm.

For further examination of the localization between proteins, particularly for our binding events, we used kernel density estimation to find the probability density of the position of one protein in the other’s reference frame. In Fig. 4C, the left panel shows the localization of one short form with respect to the other. The most common position corresponds to a bound configuration which persists for a majority of the total ∼4 *µs* in contact. When compared to the curvature plots displayed in Fig. 2, the most frequent location corresponds to a small region of *k*_1_ < 0 and *k*_2_ < 0. However, existence in this location of preferential negative total curvature is only true for one of the two proteins (*SI Appendix*, Fig. S11), serving as a possible reason for the duration of the event. In the case of the short form’s localization relative to the long form (middle panel Fig. 4C), while there exists a common bound configuration, a multitude of other positions 20 nm away are just as common. Furthermore, this bound configuration does not correspond to a state of preferential membrane curvature for either protein (*SI Appendix*, Fig. S11). Finally, no common bound configurations are observed for the case of two long forms (right panel Fig. 4C), as the uncentered protein most commonly lies along the dotted cyan lines at 10 nm and 15 nm separation. The binding conformations we observe show the capacity for curvature and direct interactions to orient proteins, as proteins tend to be anti-aligned when bound (Fig. 4B and *SI Appendix*, Fig. S9 and S10).

To further quantify these interactions, we next plot the sampling-derived free energy surfaces as a function of separation and alignment between the proteins in Fig. 5A. We estimate this free energy by treating the kernel density estimate of the probability distribution as being proportional to the Boltzmann weighted free energy, with minimum free energy set to *F* = 0 *k*_*B*_*T*. The prominent binding configuration (at about 5 nm) for two short forms corresponds to the lowest free energy, where these proteins are anti-aligned (*θ*_12_ ∼ 70°). At larger separations, a minimum at *θ*_12_ ∼ 120° becomes more prominent. For the simulation with one long and one short conformation, no significant alignment is observable in the well-defined band of minimum free energy at 20 nm separation. Additionally, there is a band of minimal free energy corresponding to a long-short bound configuration ranging from *θ*_12_ ∼45° to *θ*_12_ ∼120°, with *θ*_12_ ∼105° the global minimum within this band. At 10 nm separation, the opposing conformations experience a slight local minimum in the free energy at *θ*_12_ ∼60°. Lastly, minimal free energy regions for two long forms largely follows a band ranging from 10 nm to 20 nm separation, without any prominent angle. We note that switching the frame of reference to the other protein in the pair changes the measured values of separation and alignment slightly due to simulation precision (*SI Appendix*, Fig. S9 and S10), although this does not influence our conclusions nor significantly modify our sampling-derived free energy surfaces (*SI Appendix*, Fig. S12). Altogether, the proteins have a preference to anti-align at smaller separations, particularly when bound, where any alignment tendency decays at larger distances.

**Fig. 5.**
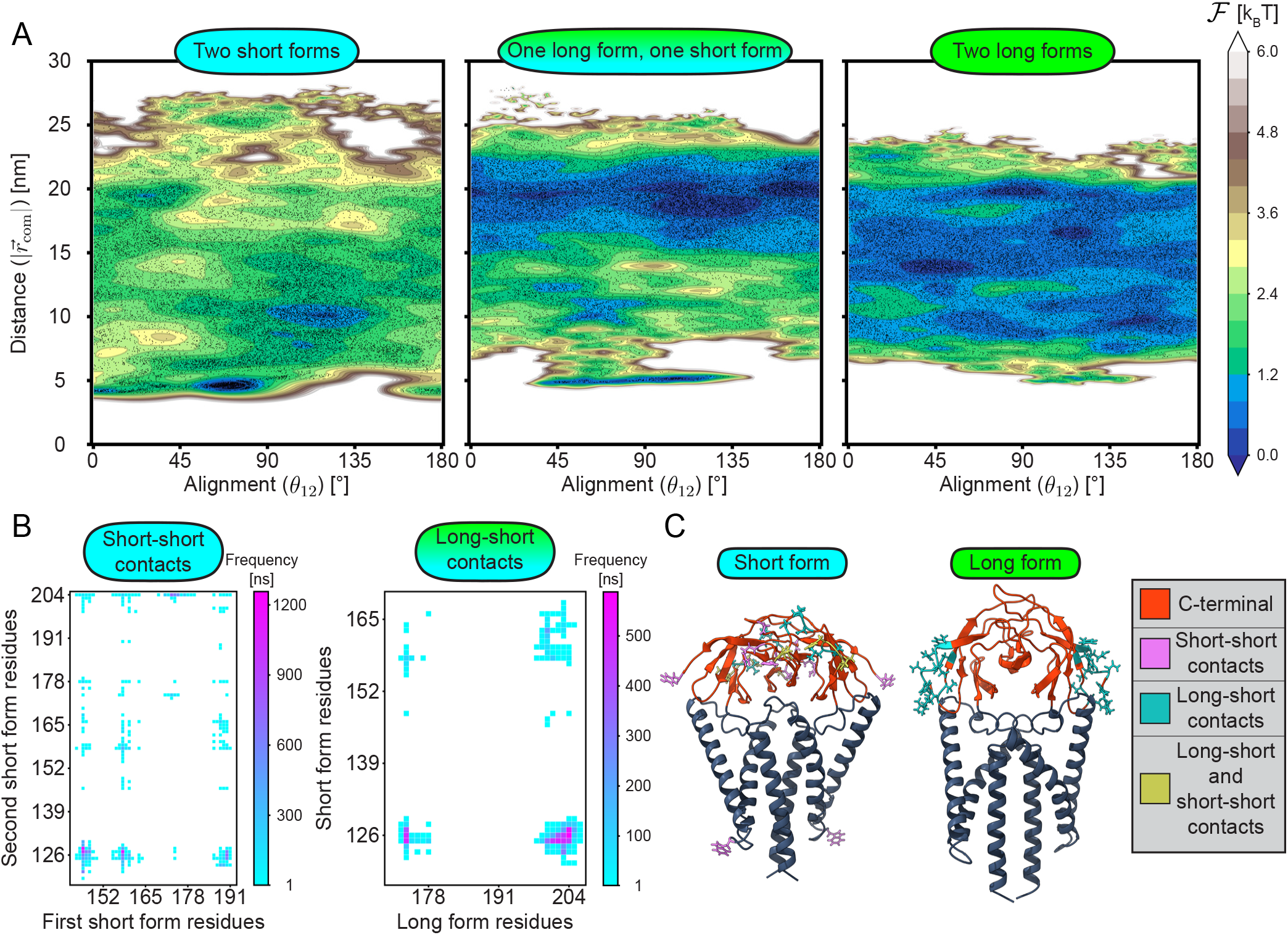
M-M interactions reflect both curvature-mediated organization and direct C-terminal contacts. (A) Sampling-derived free energy surfaces for multiple protein simulations are shown as contour plots in the separation – alignment plane, with the minimum free energy defined as *F* = 0 *k*_*B*_*T* and the plots periodic across the x-axis. Black dots correspond to data from a single simulation frame, where high free energy defines less frequent domains. (B) Zoomed in M-M contact maps for simulations with prominent contacts, where vertical and horizontal axes display residues from each individual M protein dimer. (C) All-atom short and long form structures colored according to domain and type of contact (short-short, long-short, or both). Ten of the most frequently in contact residues are colored for each contact type.

### M-M contacts are dominated by the C-terminal domain

With the vital role direct M-M interactions play in assembly and budding, we identify observed contacts during CG simulations to determine the dominant domain in these interactions. A contact between two residues from different proteins is defined to occur when the minimal distance of these residues is less than 1.5 times the size of a Martini 3 water bead (*SI Appendix*, Fig. S13). The contact frequency between residues in one dimer with residues of another is shown in Fig. 5B for two simulations with extended binding events.

Only regions with the most prominent contacts are displayed, where full maps can be seen in *SI Appendix*, Fig. S13. Identical residues in each chain of the corresponding dimer are merged into a single residue, where all residues beyond 117 are considered part of the protein’s C-terminal domain. The left-most panel shows prominent short-short contacts, with the most frequent between ARG 146 and ILE 128, for a total of 1.26 *µs* ( ∼30 % of the time bound). Furthermore, 70 % of these contacts involved C-terminal residues. Similarly, 96 % of long-short contacts were between two C-terminal residues, with the most common contact between TYR 204 and GLY 126 of the long and short forms respectively, persisting for 0.59 *µs*. Owing to the minimal number of contacts in the simulation with two long forms, no contact map is shown for long-long interactions (*SI Appendix*, Fig. S13).

Next, Fig. 5C shows the residues that are most frequently in contact for a given protein. Of the ten most prominent residues in short-short contacts, all but residue TRP 75 is in the C-terminal domain. Similarly, all prominent long-short contacts are restricted to this domain for both conformations. See *SI Appendix*, Fig. S14 for a histogram showing the frequency of these contacts for each simulation. While the residues of importance are in the long form’s C-terminal, they correspond to the edge of the protein’s major axis when looking from above (*SI Appendix*, Fig. S15). Alternatively, short form contacts are predominantly along the protein’s minor axis from above (*SI Appendix*, Fig. S15). Thus, the C-terminal domain dominates interactions between two M proteins, independent of conformation.

## Discussion

Using a combination of all-atom and coarse-grained Martini MD simulations of SARS-CoV-2 M protein dimers embedded in a membrane, we have shown that membrane curvature, which is critical for assembly and budding, is induced in a conformation dependent manner. In particular, the membrane was shown to curl around the long form’s C-terminal, forming a valley, and away from the short form’s C-terminal, forming a ridge. Thus, the long form deformation matches better with the virion bulb’s shape while the short form matches better with the neck region of a bud, in agreement with observations from SARS-CoV virions (7). While the E protein has also been shown to induce membrane curvature through MD simulations (47, 48), the sign is consistent with the neck of a budding virion. As such, these E protein induced deformations need to be very long range to be solely responsible for budding - especially when coupled with the small number of E proteins in a complete virion. Our results support the hypothesis that insufficient curvature generation is in part responsible for budding inhibition through short form stabilization (20) or the creation of a different conformation (32), as these interventions suppress long form induced membrane deformations of the correct sign.

The stark differences in curvature fields between the two conformations can be attributed to shape, as the sequence is identical. Conical proteins are known to bend the membrane away from the side with the larger domain (21, 22). The short form maintains its structure throughout our simulations, appearing conical with respect to its major axis and cylindrical along its minor axis (*SI Appendix*, Fig. S16). Solely from its shape, we would predict the membrane to bend away from the C-terminal along its major axis while hardly bending along its minor axis – in agreement with our simulated curvatures. It is to be noted that the long form shape, or cross-sections along the upper and lower leaflets, is initially like the short form. However, upon starting the simulation, a kink quickly forms in the protein’s transmembrane domain that substantially increases its cross-section along the lower leaflet with respect to both axes (*SI Appendix*, Fig. S16). This essentially switches the orientation of our cone, possibly causing the membrane to curl towards the C-terminal along both axes, as shown by long form principal curvatures. Considerable long form kink formation has been observed in other all-atom simulations (49).

We found that a single long form protein generates total curvature consistent with a budding virion, possibly driving further accumulation while creating the bud’s center. Furthermore, this localization is consistent with the long form C-terminal binding to the N protein/vRNA complex enveloped by the membrane, as observed in (7). On the other hand, short form total curvature is consistent with the edge or neck of a bud, possibly prompting its localization to this region during assembly and budding. Given the short form (33, 34) and E protein (48, 50) have also both been shown to thin the membrane, this could imply curvature and membrane thickness guide them to the edge of the bud for facilitation of membrane scission.

Our simulations of pairs of M protein conformations showed that short-short and long-long combinations tend to be closer together than the short-long pair. This is most likely a result of membrane-mediated effective repulsion stemming from the differing curvature fields between short and long forms. Furthermore, microsecond long binding events only occur in simulations involving short forms. Such binding is consistent with previous reports showing that direct interactions between short forms are sufficient for membrane clustering, an effect that is enhanced by line tension from membrane thinning (33, 34). Note that anisotropy can also lead to attraction in general (26, 27) and influence the tendency for short-short combinations to be closer together beyond binding events.

While these binding events do persist for microseconds, they are not stable, with proteins eventually separating. From the relative location of each protein in the other’s frame during these events, we can see that at least one protein is always in a non-preferential curvature region. One possibility is that insufficient direct interactions coupled to membrane fluctuations on the scale of induced membrane deformations disrupts binding over this time scale. Additionally, by examining the angle of principal protein axes, we found binding accompanies anti-alignment between proteins. Alignment or anti-alignment could be guided by curvature field anisotropy, and possibly lead to the formation of protein networks, as observed for complete SARS-CoV virions in (7). From the sampling-derived free energy surfaces for multiple protein simulations, we estimated the interaction strength between M proteins to be on the scale of ∼6 *k*_*B*_*T*, which includes thinning, curvature induction, and direct interaction contributions. Although the short form interaction energy estimated in (34) is slightly larger, this difference could result from imposed curvature or proteins never completely separating from each other. It is also possible that our sampling time may have been insufficient to find the strongest binding configuration.

We found that C-terminal domains dominated the direct interactions in M-M binding. This was particularly interesting for the long form, where the kink formation that increases its N-terminal cross-section still does not substantially contribute to binding with a short form. Experiments for the SARS-CoV-2 M protein (19) also highlight the importance of the C-terminal in higher-order oligomerization. Interactions between M proteins are most likely governed by the C-terminal at this M protein density because it extends above the membrane, whereas transmembrane or N-terminal interactions require displacement of adjacent lipids. Increasing the number of proteins could enhance non-C-terminal contacts, where proteins are forced together from direct and membrane-mediated interactions. Alternatively, particularly for long-long interactions, which we did not observe, C-terminal contacts could drive the proteins to tilt towards each other. Given the membrane curls around the long form’s C-terminal, this tilt could enhance the virion-like curvature they induce. Furthermore, the ability for C-terminal - C-terminal interactions to enhance SARS-CoV-2 budding has been shown with lower resolution coarse-grained simulations (51).

These conformation dependent behaviors provide a compelling mechanistic picture of coronavirus assembly and budding. Given the differing curvature fields for both conformations, they could naturally separate during assembly and budding, with the long form localizing and generating the middle of the bud while the short form does so for the edge. During this process the long form binds to the N protein/vRNA complex, inducing more curvature, while the short form’s anisotropic curvature field and ability to thin the membrane possibly facilitates the accumulation of E protein along the bud’s edge. The ring of short form and E protein surrounding the bud could then serve as a protein network, limiting the entry of unwanted components and driving membrane scission. As such, both conformations’ functions could not only be individually beneficial for virion formation but could co-operate with other viral proteins to enhance the process. Such a process represents a generic mechanism for viral budding even in the absence of additional host factors.

However, further confirmation that kink formation in the long form is responsible for its distinct curvature field compared to the short form is necessary to strengthen these conclusions. Additionally, while the extent in which long form curvature defines virion shape initially is clear, the observed curvature is about four times smaller than that of a fully formed virion. One possible explanation is that long form induced curvature is enhanced upon oligomerization, implying a need for the formation of a loose protein network during virion formation. Note that this requires the simulation of higher protein densities, as shown analytically in (25). These simulations with many M proteins could provide additional validation for the role of membrane-mediated interactions, identifying the coupling between localization and curvature during virion formation through comparison with discrete protein analytical models (26, 27) or mesoscopic continuum models (52, 53). It is also likely that virion curvature could rely on cooperation with N/vRNA or E protein, particularly when coupled to post-budding maturation.

Finally, while we have provided some insight into each conformation’s function, quantifying M protein conformational dynamics over the timescale of budding remains a bottleneck to our current understanding of SARS-CoV-2 assembly and budding. For example, it is possible that the short form initiates cluster formation, in agreement with our short form binding observations, and then transitions to long form. Recent studies have shown this shift is influenced by small molecules stabilizing the short form (20), while a recent preprint (54) has identified it to also be controlled by specific protein-lipid interactions. However, whether either of these transitions occur naturally during virion formation is unclear, where interactions with N/vRNA (7), high protein density, or bud curvature could also play a significant role in formational changes.

## Materials and Methods

### Molecular dynamics simulations

All-atom molecular dynamics (MD) simulations were performed using the CHARMM36m force field (55) with the GROMACS 2022.6 MD package (56) on the SDSC Expanse supercomputer (57, 58). Two different all-atom simulations were run up to 750 ns, all of which involved a 40 nm x 40 nm membrane surrounded by 0.15 M NaCl and transferable intermolecular potential with 3 points (TIP3P) water inside a periodic box. The membrane lies along the x-y plane and is normal to the z-axis, with composition identical in both leaflets and chosen to match the ERGIC: 15% cholesterol,45% 1,2-dioleoyl-sn-glycero-3-phosphocholine (DOPC), 20% 1,2-Dioleoylsn-glycero-3-phosphoethanolamine (DOPE), 7% 1,2-dioleoyl-sn-glycero-3-phospho-L-serine (DOPS), and 13% 1-palmitoyl-2-oleoyl-sn-glycero-3-phosphoinositol (POPI) (33, 34).

A single all-atom simulation was run for each embedded M protein conformation. Structures from (19) were used, where both chains of short (PDB: 7vgs) and long form (PDB: 7vgr) are referred to as a single M protein for brevity. Proteins were embedded to match previous studies (19, 31, 33, 34). Length of the periodic box along the z-axis was defined to have 5 nm of solvent between the nearest non-solvent atom and the top or bottom of the box, leading to a total height of ∼ 18 nm.

Each all-atom system was built using the CHARMM-GUI input generator (59–65). The long and short systems were prepared for simulation using the six minimization and equilibration steps provided by CHARMM-GUI. More information on these steps can be found in *SI Appendix*, Sec. 2. Post equilibration production runs were simulated in the NPT ensemble up to 750 ns with a time step of 2 fs. System temperature was maintained using the Nose-Hoover thermostat at 303.15 K (66, 67), while pressure was preserved with the Parrinello-Rahman barostat (68, 69) semiisotropically at 1 bar in the x-y plane and separately along the z-axis. Simulation coordinates were saved every 50,000 time steps (0.1 ns), for a total of 7,500 frames, with an initial frame of an all-atom simulation shown in Fig. 1A.

Coarse-grained (CG) MD simulations were run with the Martini 3 force field (44) using the GROMACS 2022.3 MD package (56). Six unique systems were simulated involving the same membrane and solvent characteristics as their all-atom counterparts: membrane only, short form, long form, two short forms, two long forms, and one long form-one short form. For simulations with two proteins, they were initially 10 nm apart according to center of mass and aligned by their principal axes. The Martinize2 script (70) was used to convert pre-minimization all-atom structures to CG, with an elastic network (71) and side-chain corrections (72) applied. To maintain tertiary structure, this elastic network was applied to the protein with a force constant of 700 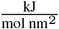, with lower and upper distance cutoff values of 0.5 nm and 0.9 nm. Note that this network is only employed within the two protein chains, allowing the monomers within the dimer to remain detached from each other.

Both membrane and solvent were created using the insane.py script (73) for each system, with a periodic box approximately 40 nm x 40 nm x 19 nm in size and a symmetric membrane composed of Martini 3 versions of 15% cholesterol (74), 45% DOPC (44), 20% DOPE (44), 7% DOPS (44), and 13% POPI (75). Surrounding solvent consisted of 0.15 M NaCl and standard water from (44). Proteins were embedded to match the all-atom systems, while also remaining similar to (19, 31, 33, 34).

Minimization, equilibration, and production files were adapted from (76), with slight modifications accounting for decreasing position restraints throughout minimization and equilibrium (*SI Appendix*, Sec. 2). After equilibration, production runs for each system were performed in the NPT ensemble up to 30 *µs*, with a time step of 20 fs. A v-rescale thermostat (77) keeps temperature at 303.15 K, while a c-rescale barostat (78) maintains pressure semiisotropically at 1 bar along the x-y plane and separately along the z-axis. For the systems involving single M proteins, a replicate with the protein restrained to its x-y center of mass was simulated for 10 *µs*. Restraints were applied through a pulling force, resulting from an Umbrella harmonic potential of strength 400 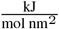.

Coordinates for each simulation are saved every 50,000 time steps or 1 ns, with Fig. 3D as an image of an initial CG simulation. Trajectory information and atom position were obtained using the MDAnalysis python package (79, 80) unless otherwise specified. Visualization of simulation trajectory frames and the generation of solvent excluded protein surfaces were performed using ChimeraX (81). Data gathering starts at 250 ns for all-atom simulations and 2 microseconds for CG versions. A summary of every simulated system is displayed in *SI Appendix*, Table S1. Lastly, the precision of every atom’s coordinate at each point in time is 0.001 nm for all-atom simulations, and 0.01 nm for CG systems. This coordinate precision contributes a small numerical uncertainty to derived quantities, particularly in two-protein analyses where reference frame transformations are compared. Nevertheless, these effects do not influence the reported trends.

### Simulation trajectory processing

Trajectories are processed either through centering or a complete transfer to the protein’s frame of reference. Both methods are done using GROMACS ‘gmx trjconv’, where centering translates the system such that the protein is translationally fixed to its initial position at every simulation frame, while keeping atoms within the periodic box. Transferring to the protein’s frame of reference involves an identical centering process followed by fitting with respect to the protein’s x-y rotation. As a result, the system is rotated in the x-y plane around the protein’s center of mass to align its principal axes with their initial configuration at each point in time. Note that with this rotational fit the membrane remains a continuous surface where lipid molecules no longer lie within the periodic box. For simulations with multiple proteins, separate processed trajectories are generated with each protein centered or as the reference.

### Protein cumulative cross-section, center of mass, and average principal axis

The set of x-y positions for all atoms in the centered protein are gathered at every time beyond 250 ns (all-atom) and 2 *µs* (CG). From the combination of these points, a concave hull defining the protein’s cumulative cross-section is generated using the alphashape python package (82) with *α* = 2. Additionally, MDAnalysis is used to find the center of mass for a centered protein, which varies slightly throughout the simulation due to trajectory precision. For systems in the protein’s reference frame, its principal axes at each point in time are calculated using MDAnalysis and are 180^*°*^ symmetric. Of the three principal axes, the first and second principal axis are predominantly in the x-y plane, and as such are the minor and major axes in the plane of the membrane. Both the time average and standard deviation are calculated using the circular mean and circular standard deviation respectively (83). When plotting, the time average of the major axis is displayed.

### Average membrane midplane height determination

To classify lipids in the upper or lower leaflet for every simulation frame of interest, the leaflet finder algorithm from (80) is applied to phospholipid head atoms with a cutoff distance of 1.4 nm. For the determination of average membrane midplane height, a 20 x 20 square grid in the x-y plane centered around the protein’s average center of mass is generated. Grid length is defined to be the maximum x-y box length experienced throughout the simulation. Note that lipid atoms were shifted into the grid during each frame after processing for the centered system, since GROMACS defines the box relative to a corner instead of the center. This was not the case for systems completely in a protein’s reference frame, with lipid atoms farther than half the box length from its center neglected to limit occurrence of bins without lipids.

The height of every phospholipid head was projected onto this grid, creating an average surface for both leaflets at each point in time. These two surfaces were then averaged together to find a grid defining midplane height, which was in turn temporally averaged throughout the simulation, leading to a grid for average membrane midplane height. For all-atom simulations, additional smoothing according to a Gaussian weight was performed on this average midplane height grid (*SI Appendix*, Sec. 1).

From this low-resolution grid, radial basis function interpolation (84) was used to generate a C2 smooth polynomial defining average membrane midplane height. The python package RBF (85) globally generated this polynomial describing the surface with a 5th order polyharmonic spline basis. Each system’s resulting polynomial minimizes the leave-one-out cross validation value, which describes how accurately the interpolating polynomial can predict a given point in its absence. In systems where the protein is only centered, this interpolation can extend between periodic images. For systems completely in the protein’s reference frame, points greater than half the x-y box length from the center are not considered. Visualization of resulting polynomials is done using the vedo python package (86), where final meshes add twenty points between every original grid point for improved visualization. Meshes of average membrane midplane height are shifted vertically such that the average height along the boundary is zero.

### Principal curvature calculation

The principal curvatures at a point along a surface are defined as the maximum (*k*_1_) and minimum curvatures (*k*_2_) experienced, where *k*_1_ > *k*_2_ with 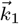 perpendicular to 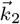 (Eq. 1) (87). A positive principal curvature corresponds to the membrane curling around the C-terminal, while negative involves curling away from the C-terminal. With a polynomial describing average membrane midplane height in the protein’s frame of reference, these quantities and their directions are determined by finding the eigenvalues and eigenvectors of the shape operator (87). This operator (S) is defined below in the Monge gauge (Eq. 2), with x and y derivatives in average membrane midplane height as h_*x*_ and h_*y*_ (*SI Appendix*, Sec. 3),

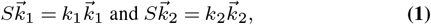

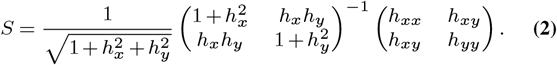

Note that we define our sign convention such that a sphere with the normal pointing inwards (as the C-terminal orients itself) has positive curvature with no additional significance (88). Plots of principal curvatures involve the same visualization process as average membrane midplane height. Principal directions are sign degenerate, and thus are modulo 180^*°*^ and added to original grid points as double-sided arrows.

Principal, and other corresponding curvatures, defined by our polynomial are taken from the four original grid points closest to the protein’s average center of mass and then averaged. This average represents the protein induced curvature, where full error estimation displayed in Fig. 3 relies on the standard deviation for these four points and the intrinsic error of each involved point (*SI Appendix*, Sec. 4). The local direction of both curvatures surrounding the protein is found through a similar process, with the circular counterpart of mean and standard deviation applied alternatively.

### Bud formation ansatz

We define a membrane cross-section during the onset of budding using a family of shapes provided from (46),

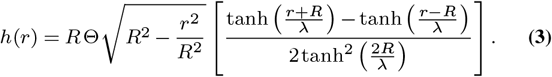

The distance from the bud’s center is *r*, where 2*R* serves as itsy width, *R*Θ its amplitude, and 2λ defines the spatial scale in which curvature switches sign and relaxes along the edge. We chose these parameters to be *R* = 65, Θ = 0.02, and λ = 125 to qualitatively match with initial bud formations.

### Separation and alignment between proteins

Protein-protein separation and alignment were calculated from the final centered trajectory for both proteins in each system, where the uncentered protein was not limited to the initial box boundaries. For each trajectory beyond 2 *µs*, both proteins’ center of mass at each time was measured using the GROMACS ‘gmx traj’ command, while their principal axes were calculated using the ‘gmx principal’ command. Separation is defined as the minimum center of mass distance between the two proteins, accounting for periodic images. Additionally, alignment between proteins is the difference between the angles (centered *θ*_1_, uncentered *θ*_2_) each protein’s major axis makes with the center of mass vector separating them 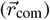. All angles are modulo 180^*°*^, where 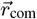 points towards the centered protein. Note that these quantities change slightly dependent on which protein is centered due to trajectory precision, serving as one form of error.

### Localization in multiple protein simulations

To find the position of an uncentered protein in the centered protein’s reference frame, the major axis of the centered protein is rotated to align with the y-axis. Major axis symmetry is ignored to prevent the centered protein from flipping back and forth, keeping a consistent monomer position. This same rotation is applied to 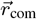, providing the localization of the uncentered protein in the centered protein’s reference frame.

With translational motion slower than rotational for the protein, estimating the localization probability distribution requires a method capable of resolving uncentered protein arcs around the centered protein. As such, a two-dimensional variable bandwidth kernel density estimate was utilized to calculate the localization probability density (89). The sample point estimator (90) from the Kernel Smoothing R package (91) was applied to the data using Python via the rpy2 Python-R bridge (92). For visualization, a 200 x 200 estimation grid was aligned with the middle protein’s center of mass, spanning 30 nm in both directions. Numerical instability in the resulting probability density is at most on the scale of 10^*−*10^ and leads to the occasional negative probability. As such, before normalizing this density with respect to the maximum, all negative values are converted to zero. This plays zero effect on the total probability, where normalized frequency in Fig. 4C begins at 10^*−*5^ to neglect these instabilities. Note that the estimate is independent of x-y bin size.

### Sampling-derived separation and alignment free energy surfaces

The same two-dimensional sample point estimator is used to determine probability density in the alignment-separation plane. Periodicity in protein alignment is accounted for by duplicating our data set twice and shifting the alignment angle by ± 180^*°*^. With this, a 600 x 600 grid is generated extending from − 180^*°*^ to 360^*°*^ in alignment and 0 nm to 30 nm in separation. For final visualization, only the middle 200 grid points for alignment and one of every three separation grid points are shown to keep consistent x-y visual resolution. The resulting sampling-derived free energy surface is simplistically defined according to the Boltzmann factor, with 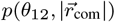 the probability distribution. Assuming a constant entropy and setting _min_ = 0 *k*_*B*_T leads to (Eq. 4),

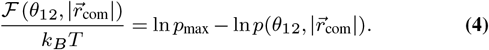

The maximum probability density observed is defined as p_max_. Probabilities less than or equal to zero, caused by numerical instabilities, were set to 10^*−*50^ for the prevention of undefined behavior when converting to a free energy.

## ACKNOWLEDGEMENTS

This work was supported by University of California Office of the President UC Multicampus Research Programs and Initiatives, grant M21PR3267 (M.E.C., R.Z, and A.G.); National Science Foundation, NSF-CREST: Center for Cellular and Biomolecular Machines at UC Merced, NSF-HRD-1547848 and NSF-HRD-2112675 (A.G.); National Institutes of Health, NIH G-RISE, T32GM141862 (J.M. and A.G.); National Science Foundation, Center for Engineering Mechanobiology, grant CMMI-1548571 (A.G.); National Science Foundation, Pinnacles Computing Cluster, NSF-ACI-2019144 (J.M.); National Science Foundation, NSF DMR-2131963 (R.Z.); and NSF–ANR (MCB/PHY) Award No. 2413062 (R.Z.). This work used Expanse GPU at SDSC through allocations BIO220146 and BIO240177 from the Advanced Cyberinfrastructure Coordination Ecosystem: Services & Support (ACCESS) program, which is supported by U.S. National Science Foundation grants #2138259, #2138286, #2138307, #2137603, and #2138296 (J.M and A. G.). The funders had no role in study design, data collection and analysis, decision to publish, or preparation of the manuscript.

---

**Fig. S1.**
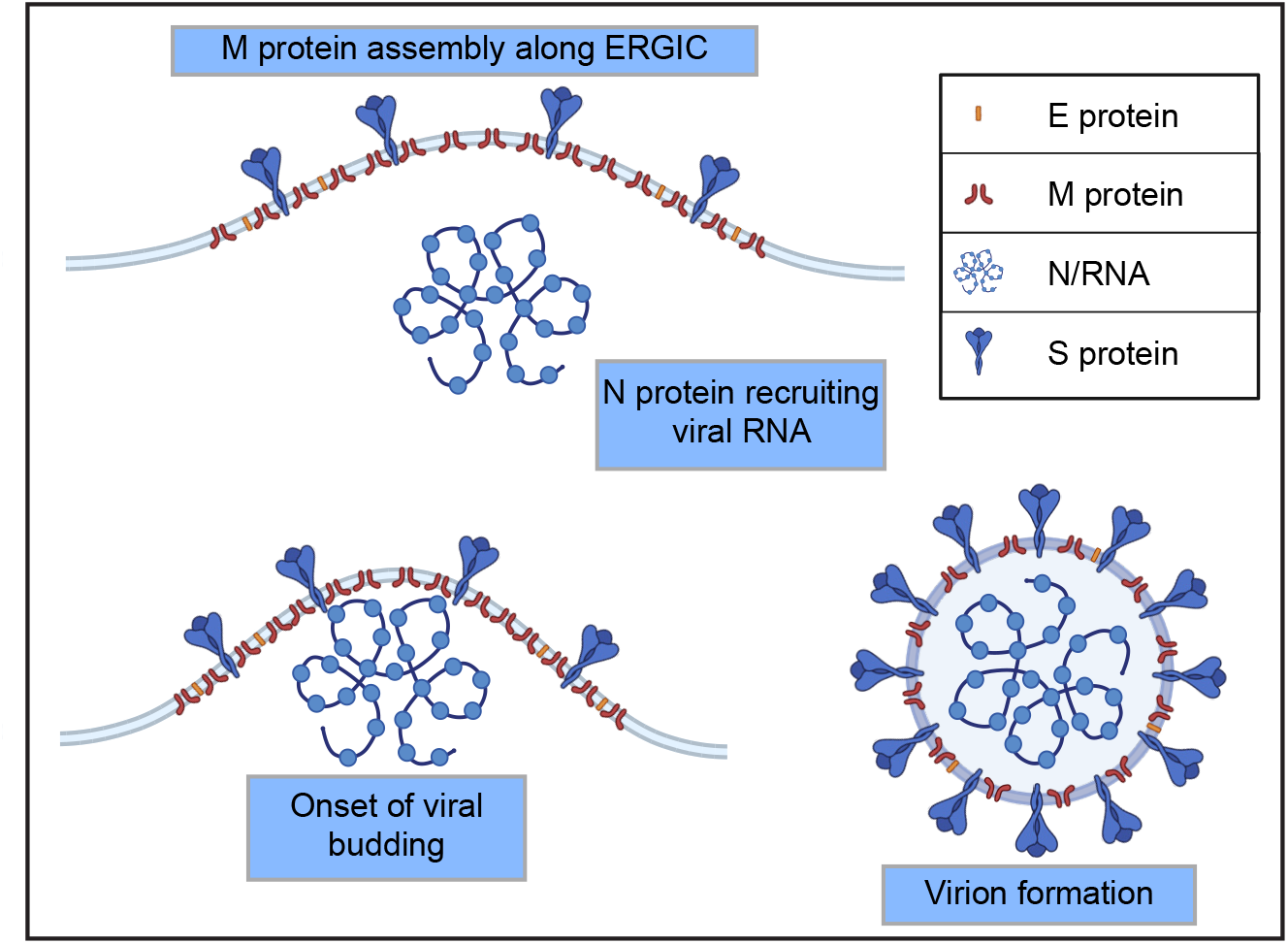
Viral assembly and budding for SARS-CoV-2 is driven by the M protein. Upon the accumulation of viral structural proteins along the ERGIC membrane, M protein guides the formation of potentially curvature inducing protein clusters occupied with M, S, and E proteins. Simultaneously, N protein recruits viral RNA, where this resulting complex is then bound to these membrane embedded clusters through interactions with the M protein’s C-terminal domain. Potentially, this further enhances curvature induction, leading to the envelopment of the virus’ genetic material and ultimately virion formation following membrane scission. Created with BioRender.com.

**Fig. S2.**
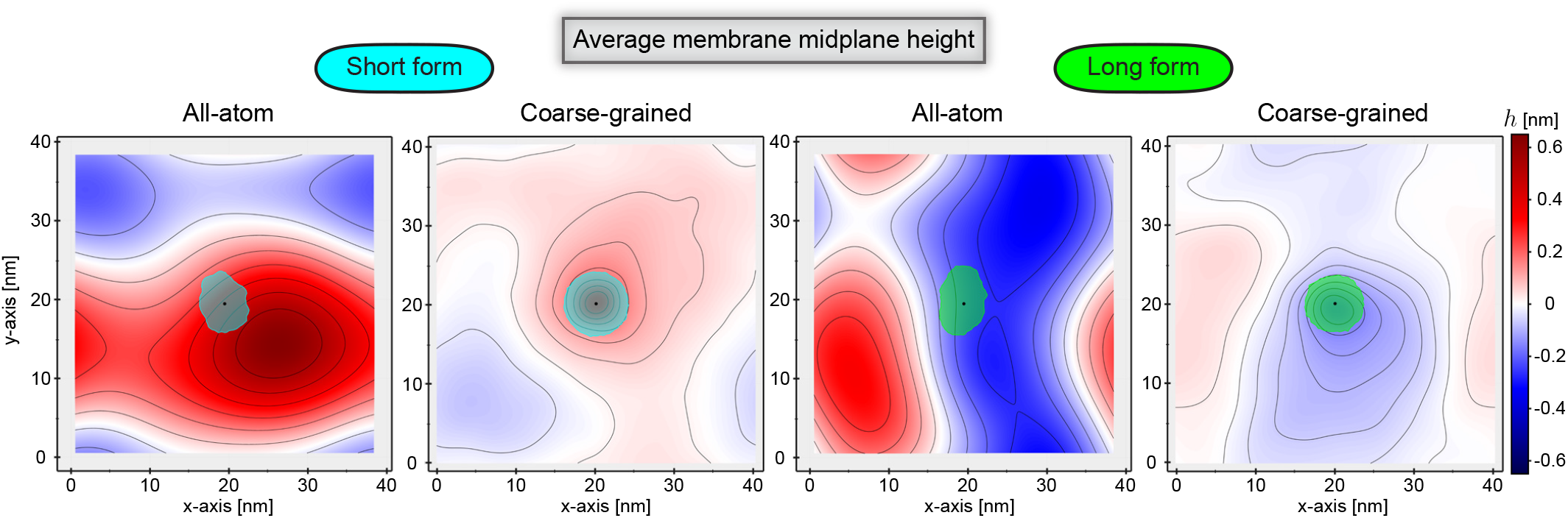
Molecular dynamics simulations of single M protein dimers, with trajectory processed such that proteins are translationally fixed to their initial position, show the short and long forms induce positive and negative deformations respectively. All-atom (250 ns to 750 ns) and coarse-grained (2 *µs* to 30 *µs*) average membrane midplane height (h) meshes are shown for both conformations - with height isolines in black. Each mesh is continuous across the boundaries and scaled equally, where boundaries differ slightly with simulation and the average h along the boundaries is set to be zero. The cumulative cross-section of the short and long forms are cyan and green respectively, with the average center of mass a black dot.

## Supplementary Text 1: Gaussian smoothing

With the comparatively small time scale explored in our all-atom simulations, we apply additional smoothing using a Gaussian weight to our raw average membrane midplane height grid. The standard deviation (*σ*_*s*_) of this Gaussian is 2.5 times the nearest neighbor distance between points, which is approximately the thickness of a lipid bilayer. For a given point, only grid points less than ten grid lengths (*dl*) away were included (Eq. 5),

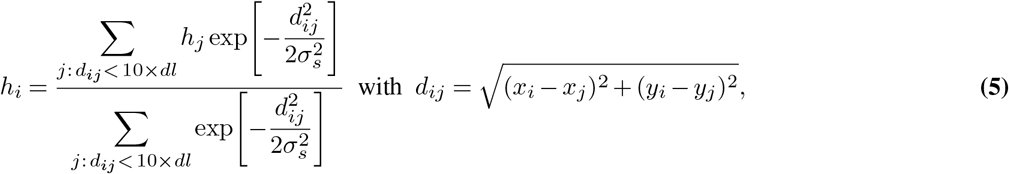

where *h* is the average membrane midplane height, with *i/j* indices corresponding to the point of interest and a single point included in the smoothing respectively. The position of the i’th grid point is *x*_*i*_ and *y*_*i*_, with the distance between the *i*’th and *j*’th grid points as *d*_*ij*_. Note that due to short simulation time, in the absence of smoothing there is too much noise between grid points to accurately quantify curvature (Fig. S3). Furthermore, the act of smoothing alters the resulting curvature magnitudes.

**Fig. S3.**
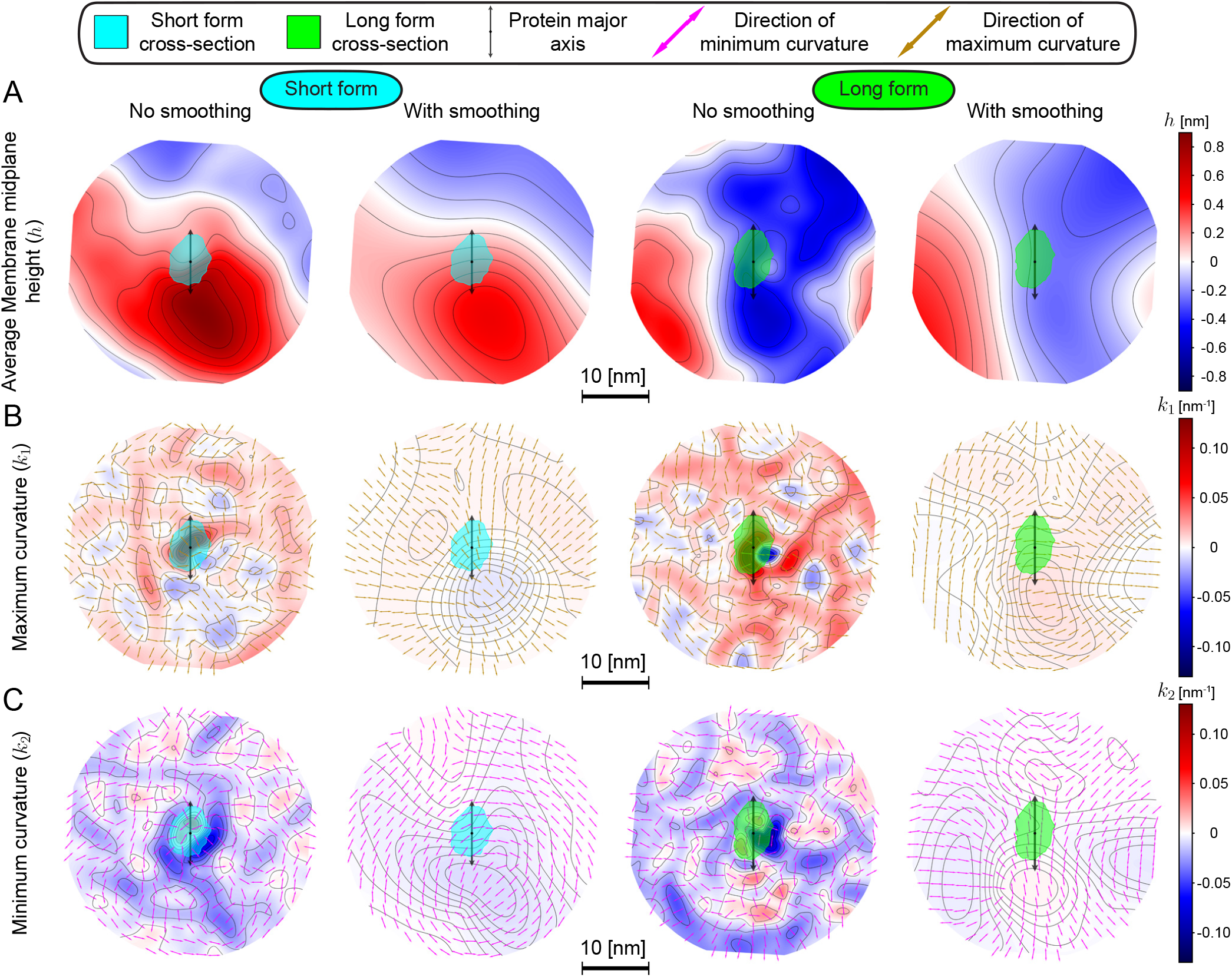
Final all-atom meshes resulting from Gaussian smoothed raw average membrane midplane height grids. Meshes of (A) average membrane midplane height (*h*), (B) maximum curvature (*k*_1_ ), and (C) minimum curvature (*k*_2_ ) are shown for both conformations, with every mesh scaled identically according to the provided 10 nm scale-bar. Black lines along the surface serve as isolines for the quantity of interest, while average height along the boundary is set to be zero. Each protein’s cumulative cross-section is colored in cyan for the short form and green for the long form, where the protein’s major axis is a black double-sided arrow oriented vertically. For each conformation, the left-most panel in (A) corresponds to a final mesh without smoothing, whereas the right-most column is the result after smoothing the original raw simulation data with a standard deviation 2.5 times the grid length. The polynomials describing these meshes are then used to find maximum and minimum curvature magnitudes and directions, with resulting meshes displayed in (B) and (C). Direction for maximum and minimum curvatures are shown as brown and pink double-sided arrows respectively. Thus, all-atom simulations are used primarily to verify the deformation’s sign and qualitative shape. Smoothed all-atom height with a different colorbar is also portrayed in Fig. 1C.

**Fig. S4.**
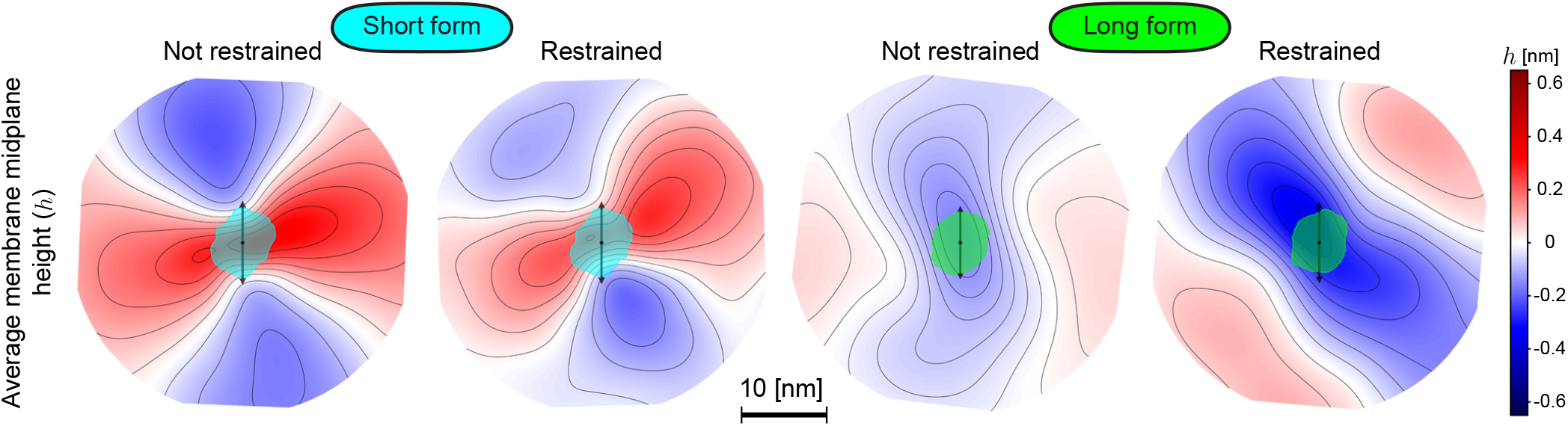
Restraining M protein to maintain its initial x-y center of mass throughout the simulation does not substantially change induced deformation. Final average membrane midplane height meshes from individual protein CG simulations are displayed for both conformations, where each conformation’s left- and right-most panels correspond to height in the absence and with restraints respectively. Meshes are generated with data from 2 *µs* to 30 *µs* (unrestrained proteins) and 2 *µs* to 10 *µs* (restrained proteins). Isolines for height are shown in black, with meshes scaled equally and zero set to be the average along the boundaries. The short form’s cumulative cross-section is in cyan while the long form’s in green, with the protein’s average major axis (black double-sided arrow) oriented vertically. For the long form, restraints minimally increase the magnitude of the deformation while slightly shifting the bend in the membrane away from its major axis. Alternatively, restraining the short form barely reduces the deformation it induces without altering the orientation of the bend. Since restraining each protein prevents lateral diffusion to pre-existing membrane curvature extrema, the lack of substantial curvature field alteration supports curvature induction over or in addition to curvature sensing. Unrestrained meshes, or the first and third columns, are also displayed in Fig. 1C.

**Fig. S5.**
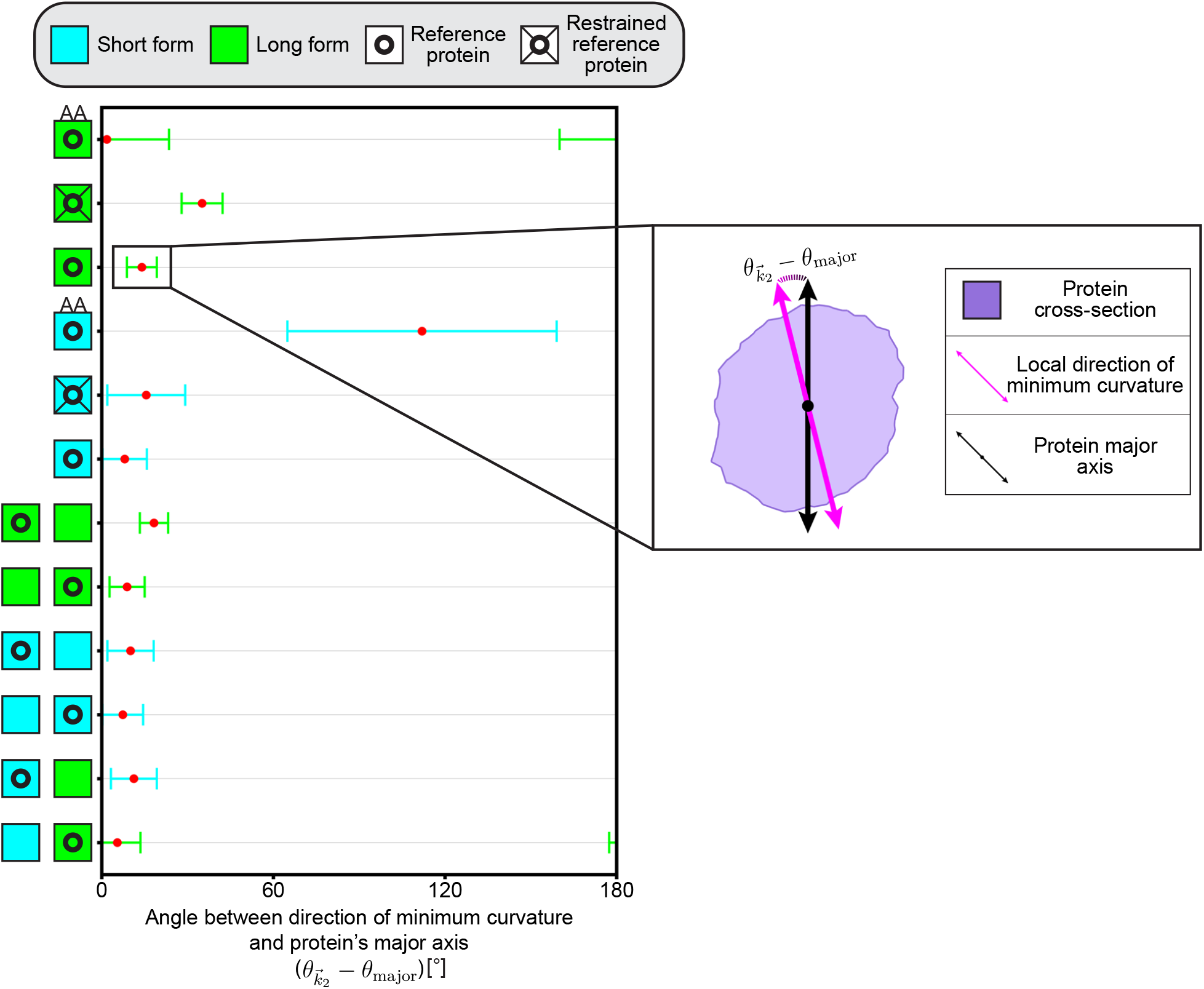
Nearly every M protein conformation’s major axis is aligned with the local direction of minimum curvature. The angle between local 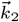 and a protein’s major axis is defined as 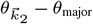, where 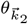 is found by averaging this direction adjacent to the protein’s center of mass. Note that every angle is modulo 180^*°*^. This quantity is periodic across the x-axis of the provided chart and is shown with respect to each protein’s reference frame for all simulations, with simulations labeled according to type and number of proteins by colored squares along the y-axis. A green square represents a long form whereas a cyan square signifies a short form - a system with two proteins will have two squares. Reference proteins, or the protein 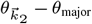 is being measured for, are labeled with a single circle inside a square. Observed error is shown as x-axis spread around the determined 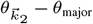 (red dot), and is colored according to the reference protein. All-atom results are included with an additional ‘AA’ label on top of the square, while restrained proteins are depicted with lines connecting the circle to the surrounding square. An example schematic showing the orientation of local minimum curvature (pink double-sided arrow) and the protein’s major axis (black double-sided arrow) for the single long form CG simulation is shown in the inset on the right, with the cumulative cross-section of the protein purple. Other than the all-atom short form and restrained CG long form simulations,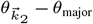 is within error and close to zero for every simulation.

**Fig. S6.**
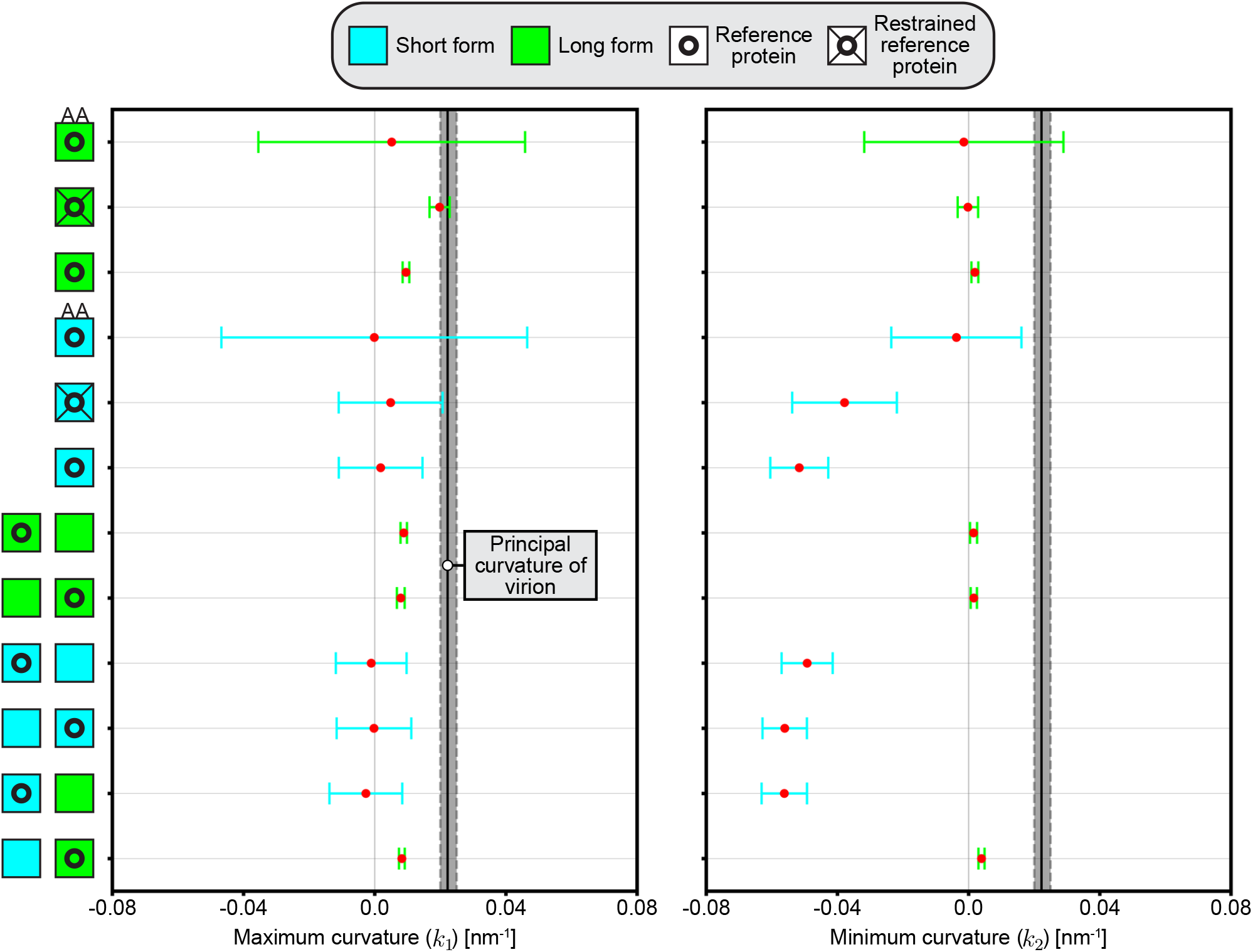
The long form induces positive maximum (*k*_1_ ) and minimum (*k*_2_ ) curvature, while the short form induces an anisotropic ridge with approximately zero maximum and negative minimum curvature. Both *k*_1_ and *k*_2_ are found through a local average surrounding each protein’s center of mass, with all simulations labeled according to the type and number of proteins along the y-axis. Green and cyan squares correspond to long and short forms respectively, where two squares implies a system with two proteins. The principal curvature for a given reference protein, identified by a circle inside a square, is shown as a red dot with error as x-axis spread colored according to protein. Restrained proteins are signified with lines connecting the circle to the surrounding square, while all-atom simulations are labeled with ‘AA’. Both charts include the virion’s principal curvature,or 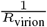 as a vertical line for reference, with *R*_virion_ = 45 nm ± 5 nm. Note that outside of the all-atom simulations with pronounced error due to shorter simulation time and the slightly altered principal curvature magnitudes for restrained proteins, each long and short form induces a consistent principal curvature within error. Short form error is more pronounced due to its spatially refined ridge formation, leading to greater curvature variation surrounding the center of mass.

**Fig. S7.**
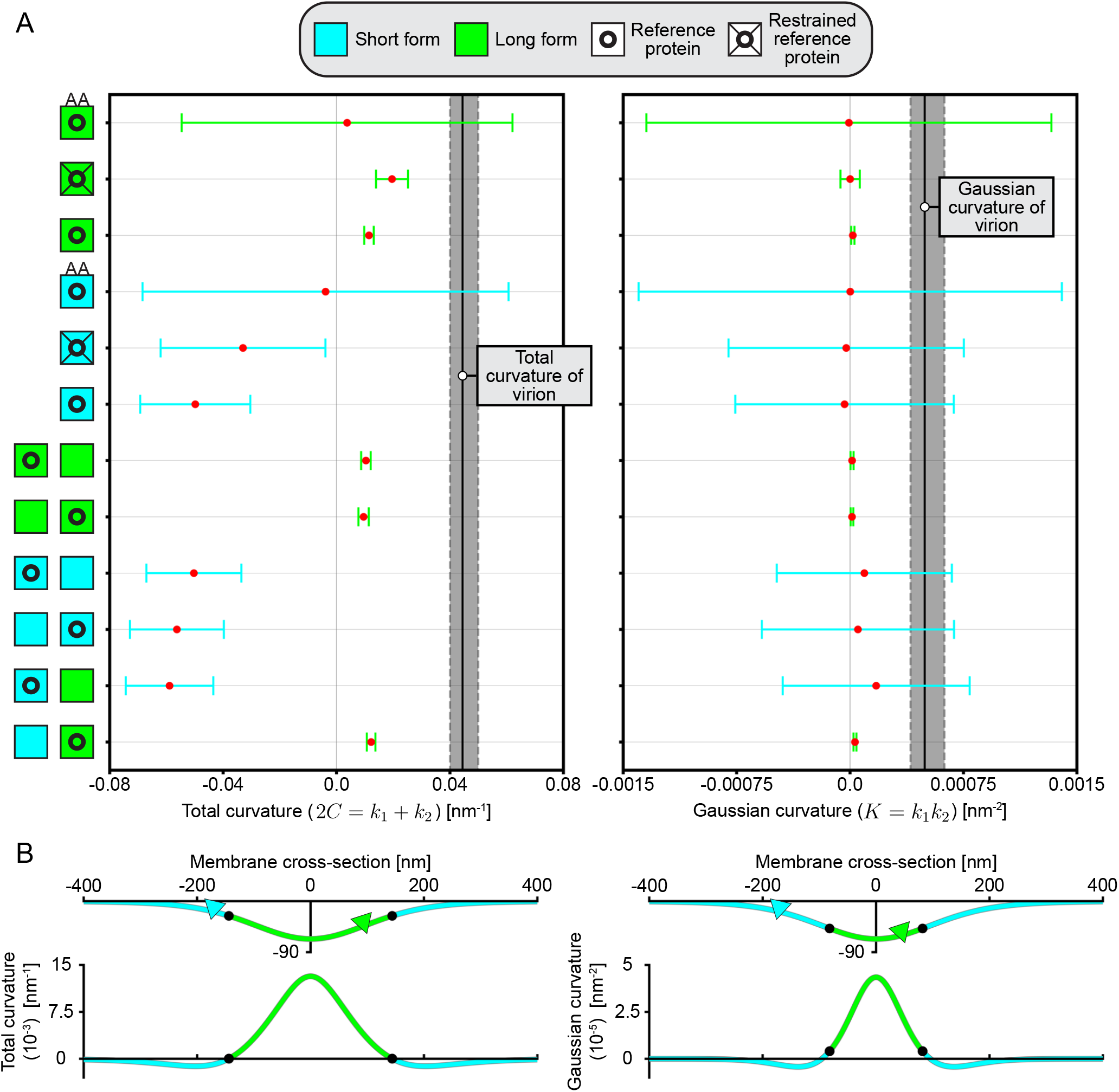
Long form induced total and Gaussian curvature is consistent with the bulb of a budding virion while that of the short form fits with the edge. (A) Charts of total (2*C* = *k*_1_ + *k*_2_ ) and (B) Gaussian (K = *k*_1_ *k*_2_ ) curvature for every protein in each simulation. Both quantities are found using average *k*_1_ + *k*_2_ and *k*_1_ *k*_2_ surrounding the protein’s center of mass. Following our previous convention, y-axis labels describe the system of interest, with green and cyan squares as the long and short forms respectively. The measured curvature (red dot) is induced by the reference protein (circle inside a square), with x-axis spread the error (colored by protein). A system with two squares has two proteins, yet only a single protein is ever the reference. Total and Gaussian curvature of the virion is displayed as a filled, vertical black line given *R*_virion_ = 45 nm ± 5 nm. Outside of the all-atom (‘AA’) systems, every total curvature in (A) is shown in Fig. 3A and 3C. All-atom curvature error is much more pronounced due to low simulation time and additional smoothing. Total curvature of the long and short forms is consistently positive and negative respectively. As expected given our induced maximum and minimum curvatures, the long form Gaussian curvature is always positive, whereas the short form hovers around zero given error. (B) Example membrane cross-sections during the onset of budding, showing which regions agree with long and short form total curvature (left) and Gaussian curvature (right). Green and cyan coloration corresponds to agreement with long and short form curvatures, while triangles indicate protein orientation with the C-terminal the base. Below each cross-section is the corresponding total and Gaussian curvature, displaying the chosen classification for long and short preferential regions, with black dots the transition points. While the transition points change slightly for different curvature classification methods, long and short forms consistently match with the bulb and edge of a budding virion.

**Fig. S8.**
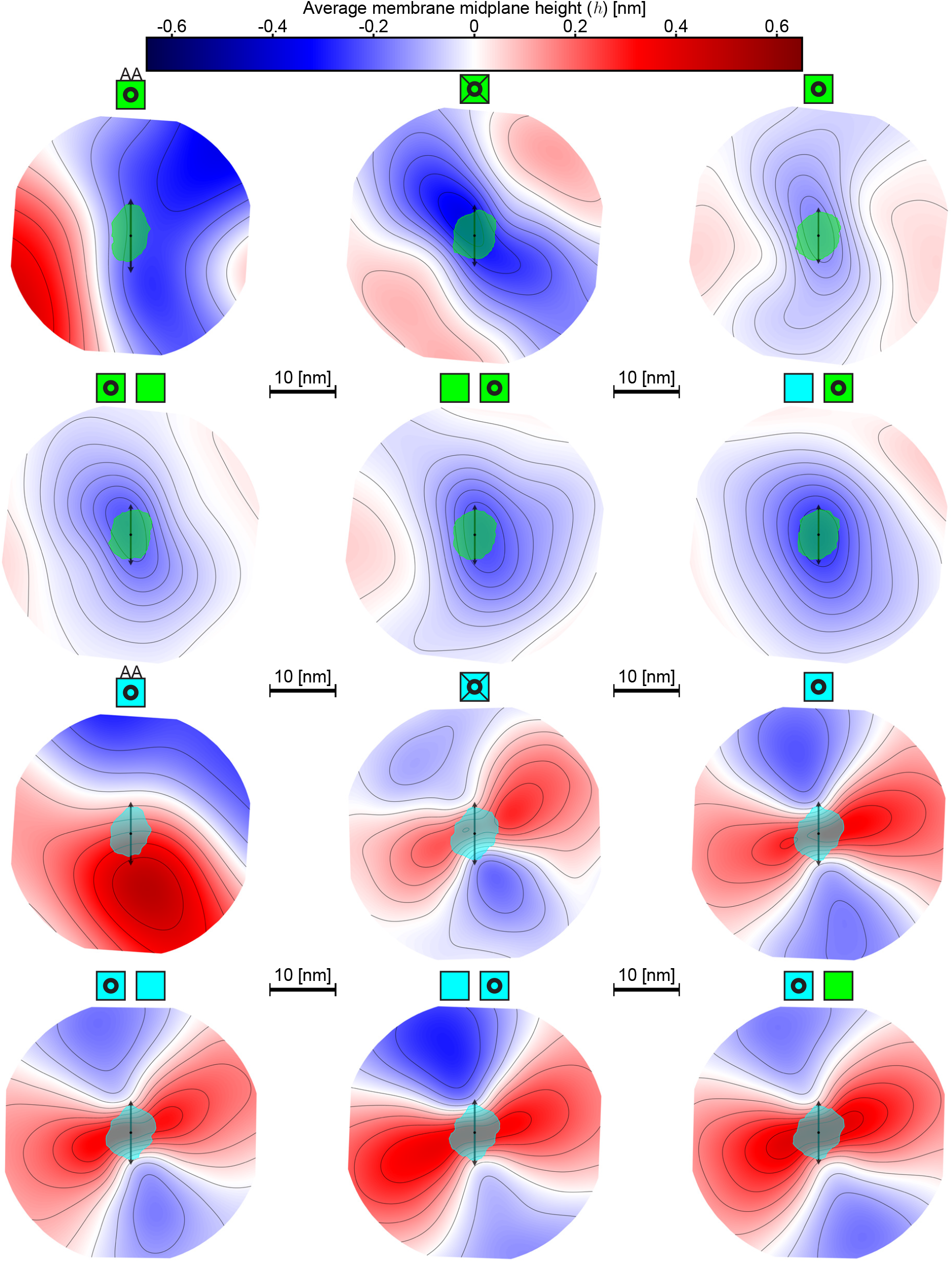
Average membrane midplane height (*h*) meshes with every protein as a reference show consistent conformation dependent deformation sign in all systems, with symmetry prominent in coarse-grained simulations. Zero is set as the average along the boundary, while isolines are displayed in black. Each equally scaled mesh is labeled according to the simulation and reference protein following our previous convention, with long (green) and short (cyan) forms indicated with a square. The reference protein for a mesh is represented with a circle inside this square, where the mesh is oriented such that the protein’s major axis (black double-sided arrow) is pointing upwards, and its cumulative cross-section is displayed in green (long form) or cyan (short form). All-atom simulations are noted with the ‘AA’ symbol, while restrained proteins are given additional lines connecting the circle to the square. Each non-restrained single protein mesh is also displayed in Fig. 1C, while restrained meshes are additionally shown in Fig. S4.

**Fig. S9.**
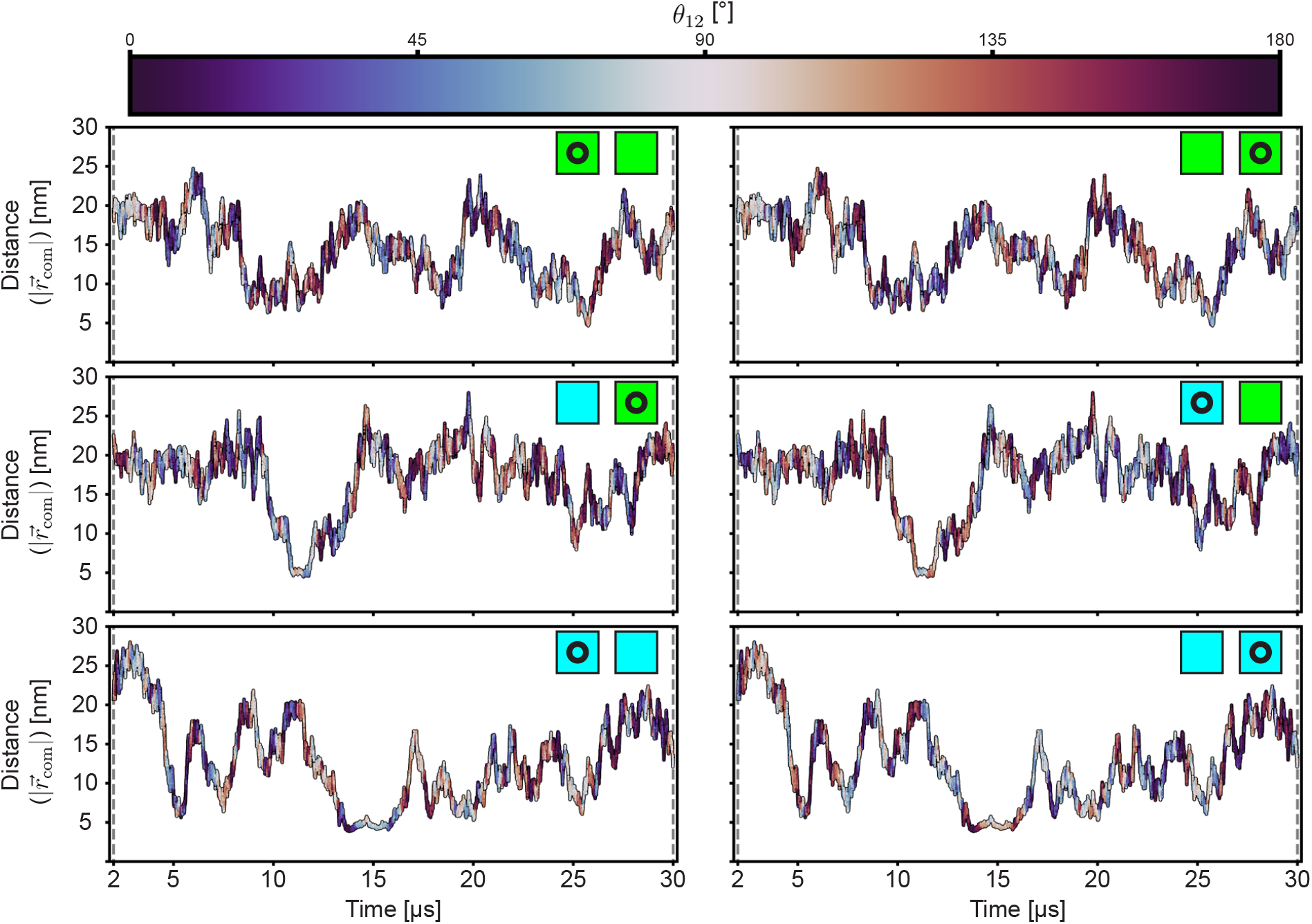
Dissimilar conformations repel, while similar ones tend to remain closer, with simulations involving a short form experiencing binding between conformations and anti-alignment. For all two protein simulations, distance 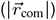 between their centers of mass is displayed at each time beyond 2 *µs*, with squares serving as labels for long (green) and short (cyan) forms. The colorbar defines alignment between both proteins’ major axis (*θ*_12_ *θ*_1_ *θ*_2_ ) at each time, where switching centered proteins mirrors the angle and all angles are modulo 180^*°*^. Each row corresponds to a single simulation, where the left and right columns differentiate centered proteins (indicated with a circle inside square protein labels). Note that *θ*_12_ is continuous in time, while *θ*_1_ and *θ*_2_ individually are not by definition due to our periodic images. Beyond mirrored alignment, variations in results between centered proteins are insignificant and solely due to simulation precision. For each of our binding events, *θ*_12_ is predominantly between ∼ 60^*°*^ and ∼ 120^*°*^, indicating anti-alignment. All three plots in the left-most column are also displayed in Fig. 4B.

**Fig. S10.**
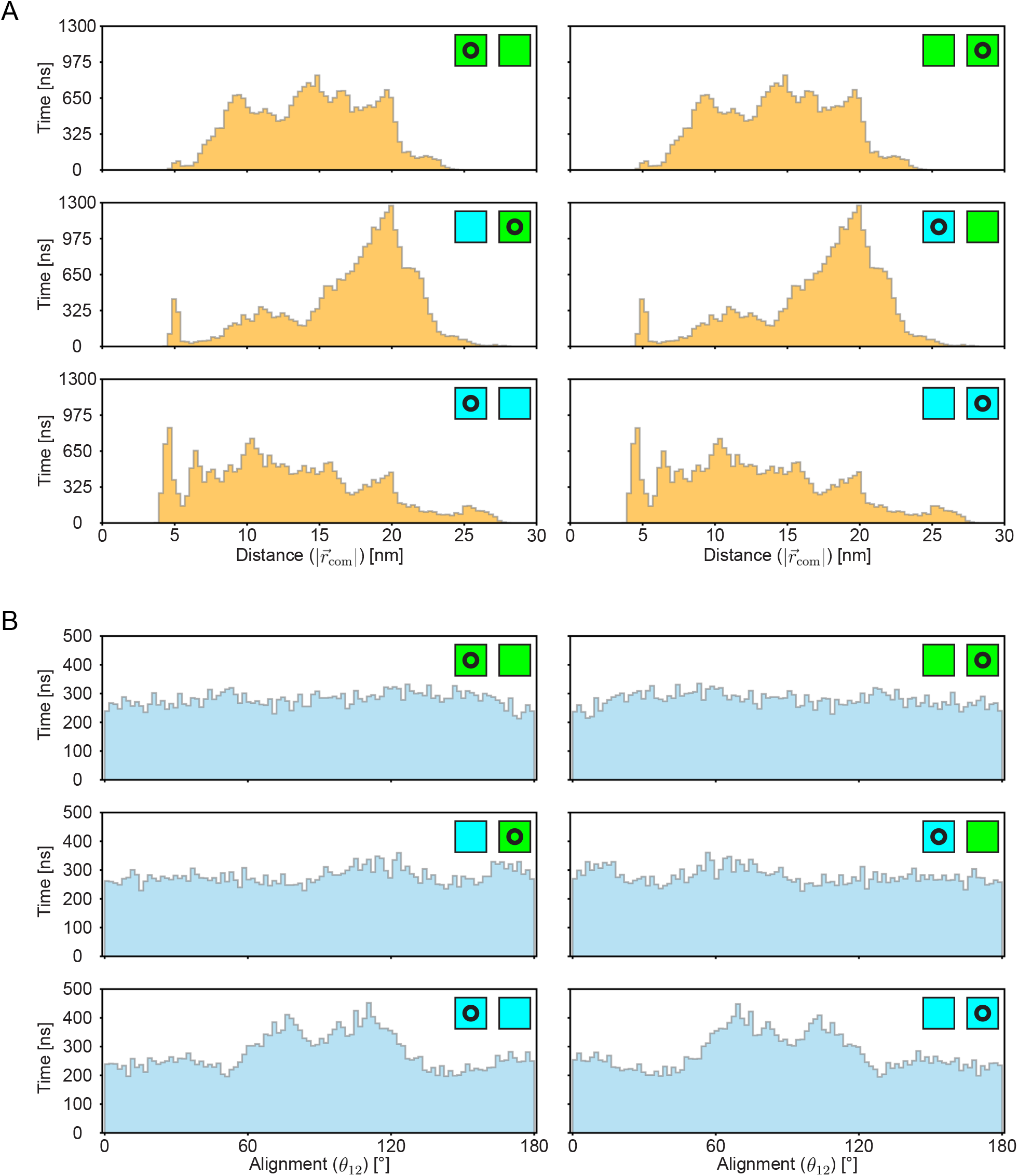
Separation and alignment histograms highlight that dissimilar conformations tend to be farther away then similar ones, while pronounced binding coincides with anti-alignment. Each histogram displays the cumulative time after binning according to (A) center of mass distance between proteins 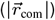, and (B) the alignment between major axes *θ*_12_ . Each row is a different simulation, labeled according to the conformation combination as green (long form) or cyan (short form) squares. Columns within a row represent a switch from one protein being centered (circle inside square) to the other. Switching which protein is centered slightly alters distance due to simulation precision while mirroring alignment. From (A), the simulation with two long forms results in a band of common distances from ∼ 10 nm to ∼ 20 nm, with the two short form simulation biased towards a distance of ∼ 5 nm. Alternatively, our system with different conformations is dramatically centered around ∼ 20 nm. Alignment histograms in (B) identify no prominent overall angles for the two long form simulation, however, our two short form system displays a clear tendency to anti-align due to the prolonged binding event. No noticeable trend appears for the simulation with differing conformations beyond a possible bump at alignments of ∼ 120^*°*^ (left) or ∼ 60^*°*^ (right). Note that this analysis washes away any coupling between 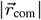 and *θ*_12_ .

**Fig. S11.**
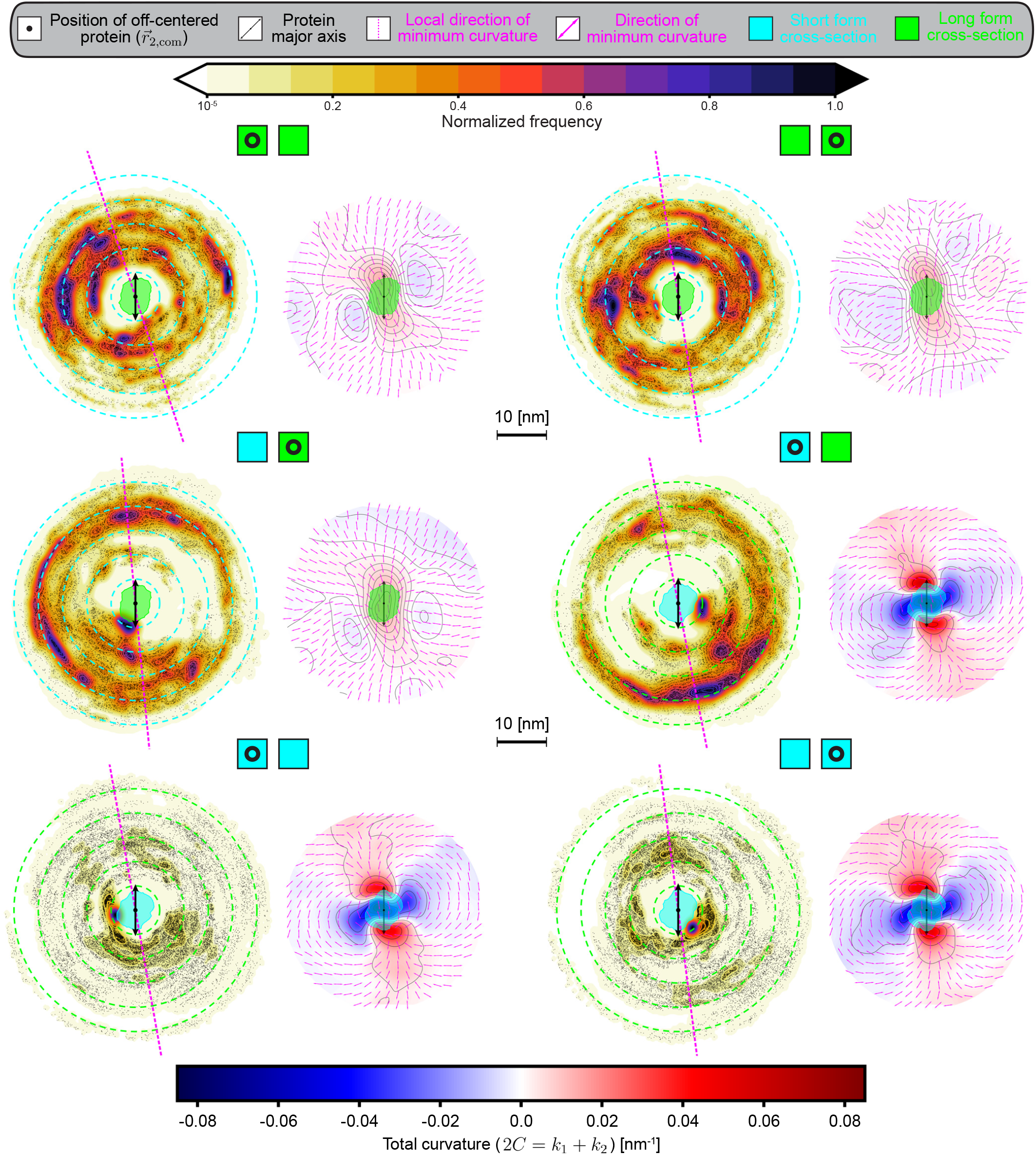
The localization of one M protein in the other’s reference frame rarely coincides with preferential curvature. Probability density for each protein in the reference frame of the other is displayed and colored according to a normalized frequency for all two protein simulations, with the location of the off-centered protein 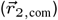 at each time represented as black dots. Protein cumulative cross-sections are colored in green (long form) and cyan (short form), with the protein’s major axis oriented vertically (black double-sided arrow). Each pink dotted line is the local direction of minimum curvature adjacent to the reference protein, while surrounding circular dotted lines identify radial distance from the center of mass in units of 5 nm. For each probability density, total curvature (2*C* = *k*_1_ +*k*_2_ ) induced by the reference protein is shown on the right as an identically scaled mesh, with pink double sided arrows portraying the direction of minimum curvature and isolines in black. Each row represents a different simulation, as indicated with the colored squares, where a circle within a square identifies the reference protein which alternates with column. Note that the left column’s probability density is also displayed in Fig. 4C.

**Fig. S12.**
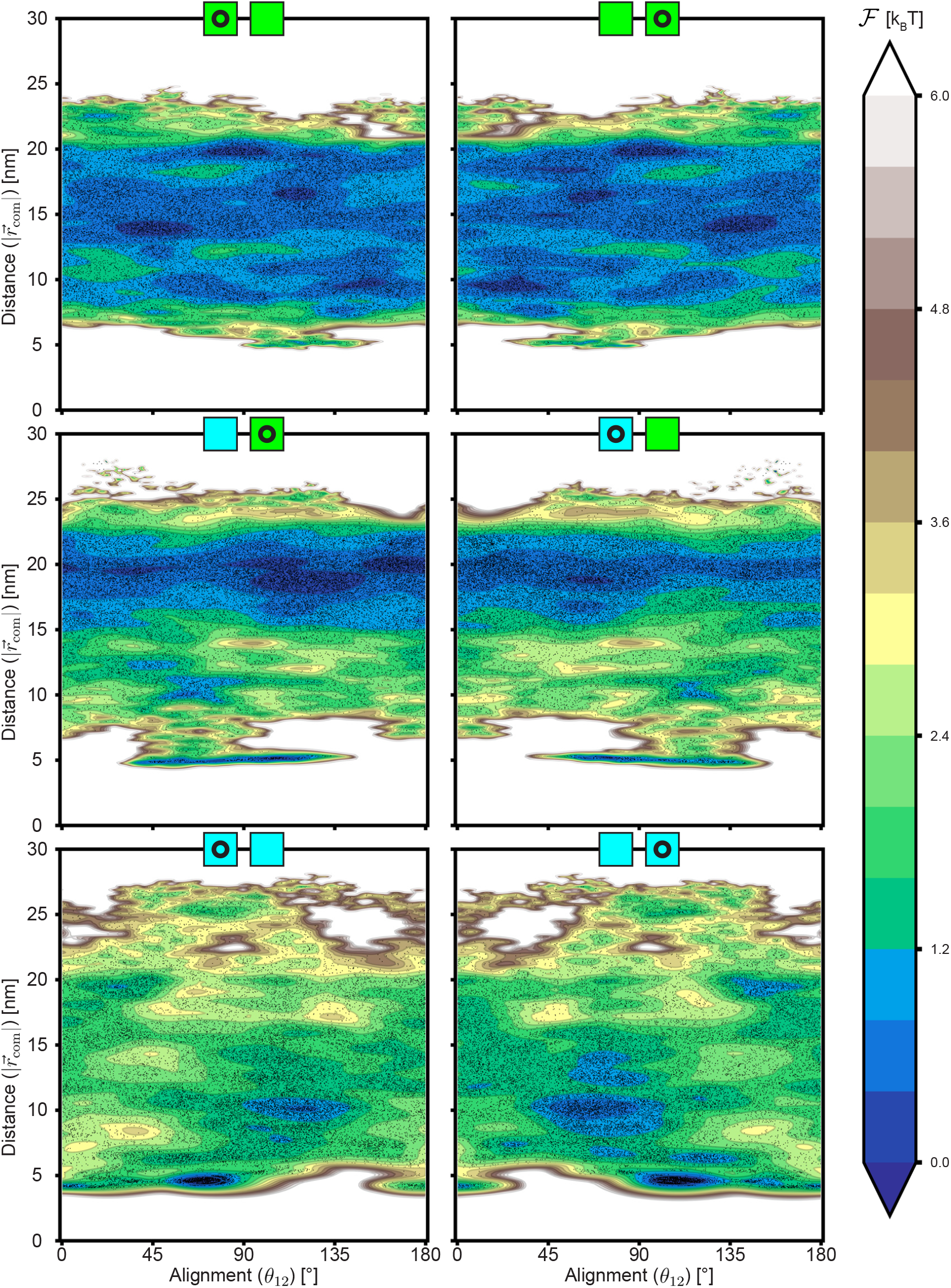
Simulation precision impacts sampling-derived free energy surfaces, but does not influence our conclusions. Separation and alignment free energy surfaces are displayed as contour plots with both proteins centered for each simulation, where the minimum free energy is set to *F* = 0 *k*_*B*_*T* . Each row corresponds to a single simulation, with squares identifying number of long (green) and short (cyan) forms, while columns within switch the centered protein (circle surrounded by square). Black dots represent the location of a single simulation frame in this plane, where every surface is periodic in alignment. All free energy surfaces in the left column are additionally included in Fig. 5A. The difference between our surfaces upon switching the centered protein, barring alignment mirroring, quantifies error from our methodology and simulation precision.

**Fig. S13.**
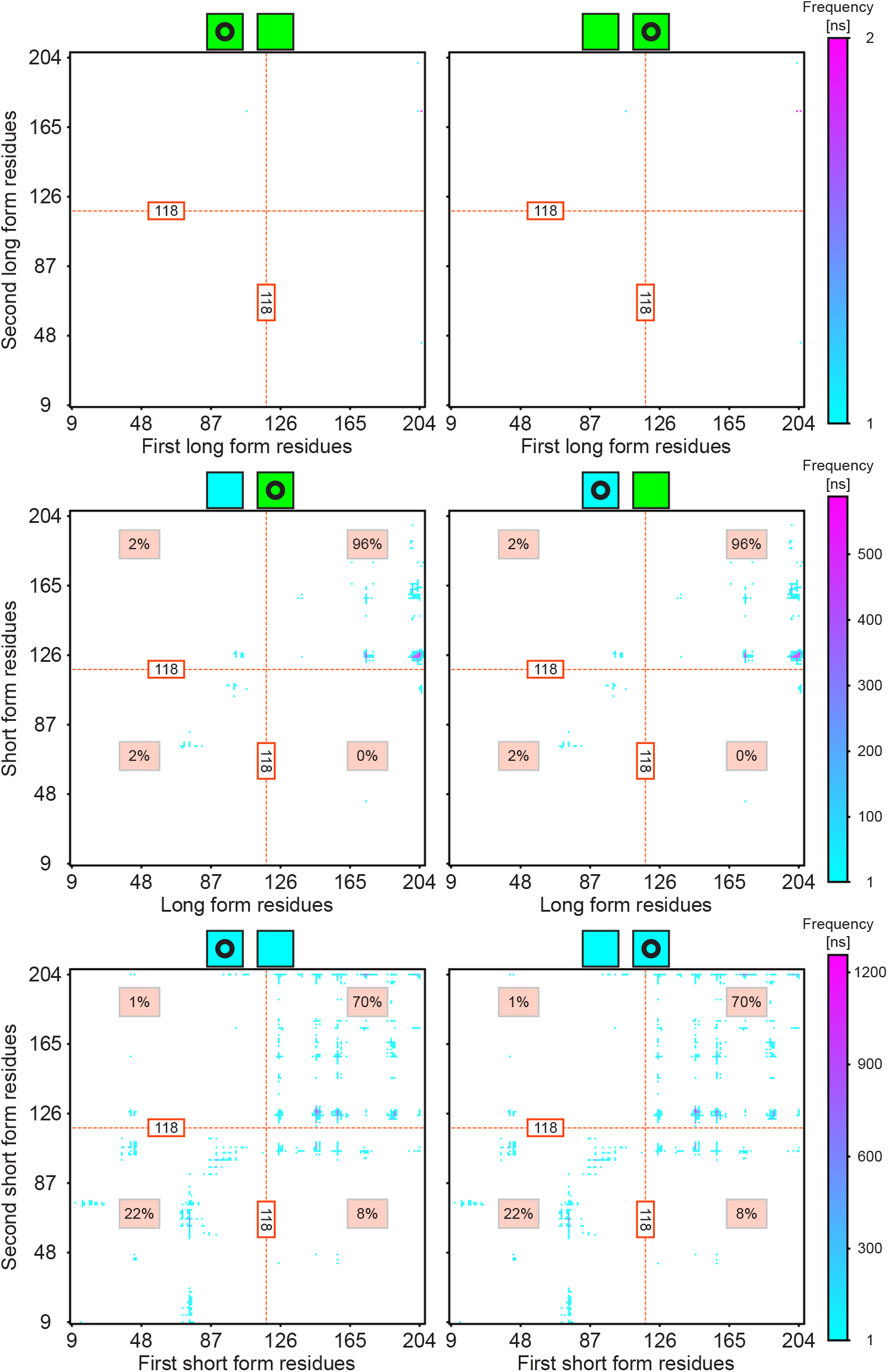
Contacts between M protein conformations are dominated by the C-terminal domain. Frequency of residue-residue contacts between two M proteins is shown for each simulation and centered protein combination. Two residues from different proteins are in contact if they have a minimum distance less than 1.5 times the size of a Martini 3 water bead (taken to be 1.5 × 0.47 nm). When counting contacts, both chains in a conformation are merged into one, leading to residue ranges of 9-204 and 9-206 for the short and long forms respectively. Residue 118 signifies the beginning of the C-terminal domain (dotted orange line), and splits maps into four different quadrants. Contacts within these quadrants then make up a percent of the total contacts as indicated. Maps in the same row are from the same simulation, with protein combinations labeled according to colored squares, and the centered protein switching with column (circle inside square). Note that axes remain the same with centered protein switching, where contacts change insignificantly with different centered proteins due to simulation precision. Zoomed in regions from the bottom two contact maps in the left column are in Fig. 5B. More data is needed to further examine long-long contacts, as the most common contact only occurs for 2 ns cumulatively.

**Fig. S14.**
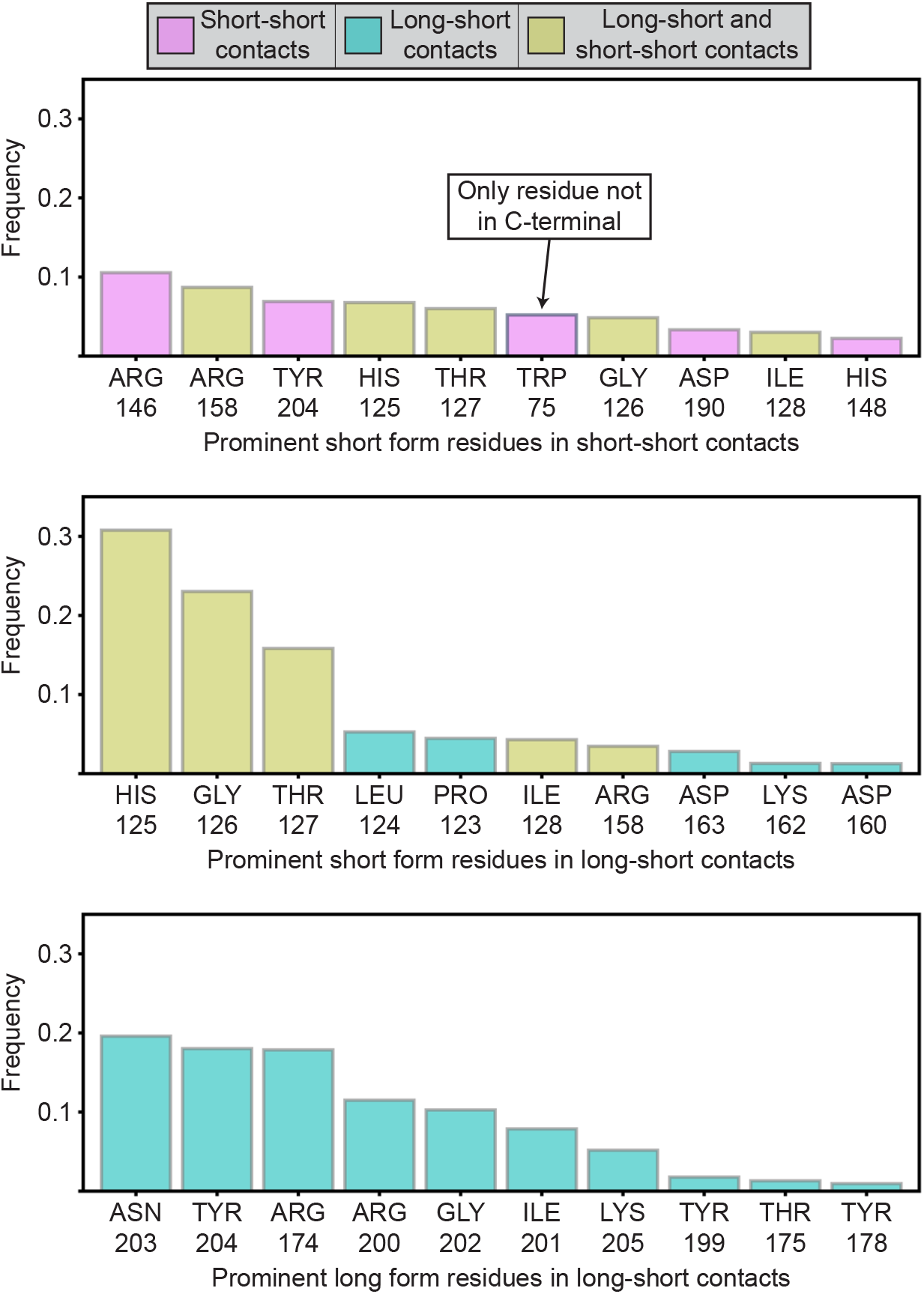
Prominent residues in short-short and long-short contacts nearly exclusively reside in the C-terminal domain. Frequency is quantified as the number of times a residue in a protein is involved in any contact divided by the total number of contacts said protein experienced. For two identical proteins, we merge all contact residues into a single pool to generate this frequency (where the total number of contacts is effectively doubled). Residues are colored according to simulation or type of contact: short-short (pink), long-short (cyan), and both (tan). The ten most prominent short form residues in short-short interactions are shown at the top. Similarly, the middle portrays the ten most prominent short form residues in long-short contacts, while the bottom does so for long form residues in long-short contacts.

**Fig. S15.**
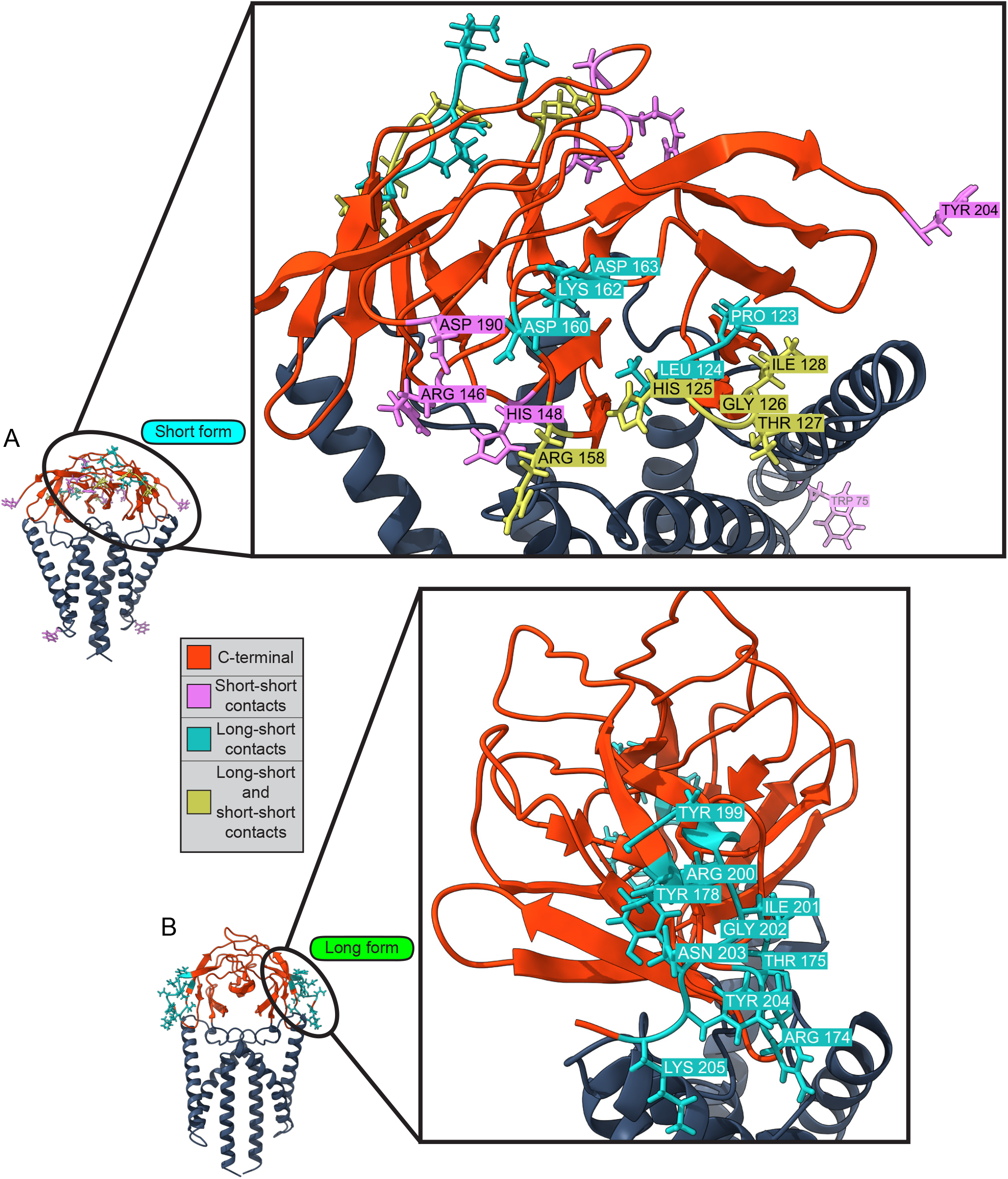
While prominent long form contact residues tend to be along the edge of its major axis in the C-terminal when looking from above, those of the short form lie closer to the edge of its minor axis. All-atom structures of the (A) short and (B) long forms are colored according to domain and type of contact: short-short (pink), long-short (cyan), and both (tan). Both insets show the exact location and name of every important residue. Left-most images are the same as those in Fig. 5C.

**Fig. S16.**
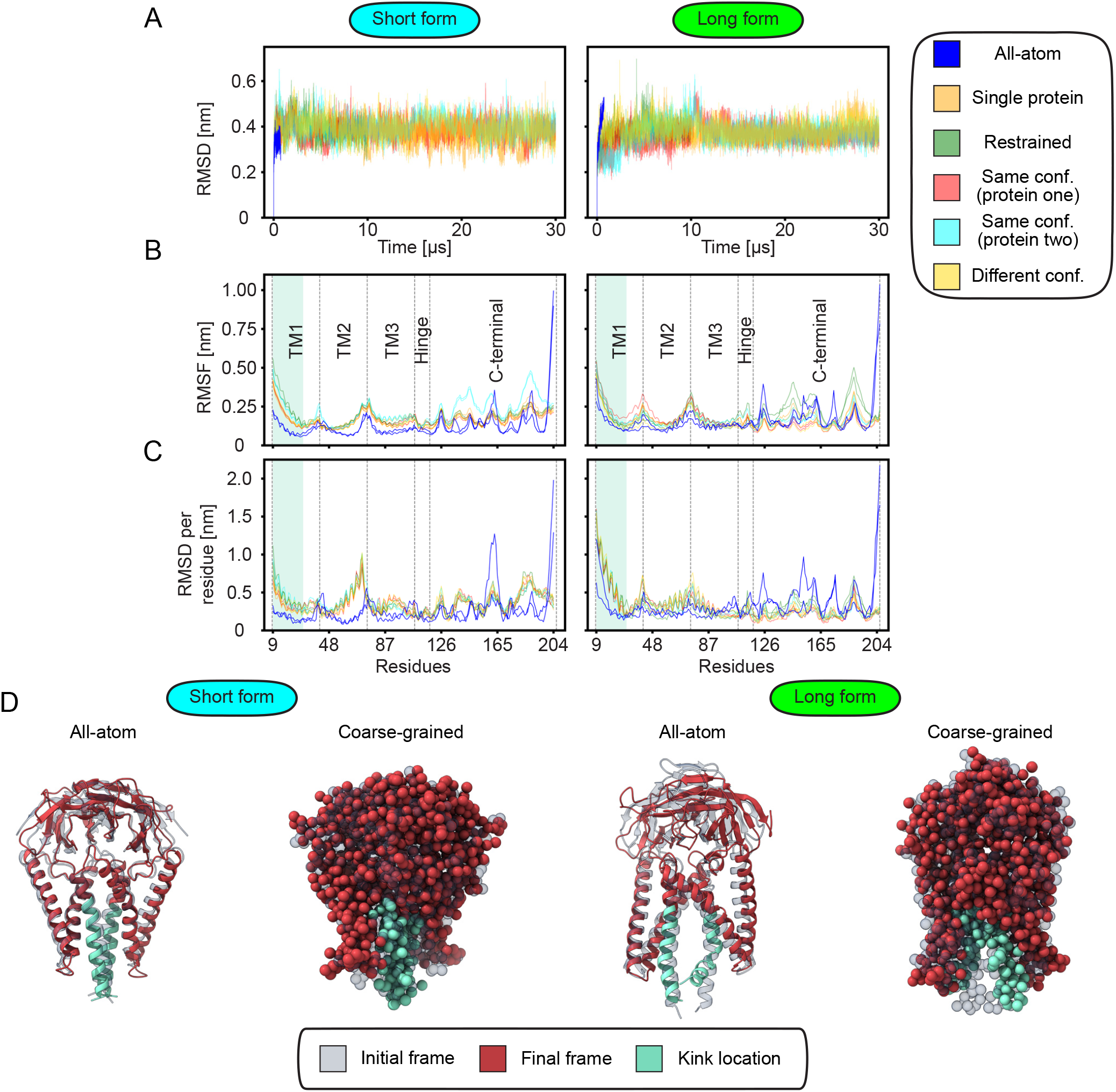
The short form maintains its structure during all-atom and coarse-grained MD simulations, while the long form develops a kink in its transmembrane domain at both resolutions. Stability is measured for each simulated short (left column) and long (right column) form, with curves colored according to simulation. All-atom results are opaque and shown in blue, while CG ones are transparent to better visualize overlapping regions. For simulations with two of the same conformation, protein one is the first dimer in the structural file, whereas protein two is the second. (A) Root mean square deviation (RMSD) serves as a quantification of structural deviation for an entire structure with respect to the initial simulation frame at each point in time. While RMSD remains within the range expected from thermal fluctuations for the all-atom short form, the long form experiences a substantial deviation from its initial structure, indicating kink formation. Since the applied elastic network has the same force constant for each protein, RMSD for Martini CG simulations is similar for every protein, but is larger and smaller than the all-atom counterparts of the short and long forms respectively. (B) Per-residue root mean square fluctuation (RMSF) defines the standard deviation of a residue about its average position, with gray dotted lines indicating different regions of the protein, and each chain within a dimer colored identically. Residues defining these dotted lines are taken from (19) as transmembrane helix one (TM1, residues ∼ 9-41), TM2 (residues ∼ 42-74), TM3 (residues ∼ 75-107), hinge region (residues 108-117), and C-terminal domain (residues ∼ 118-206). Residues 9-30 are shaded cyan, and represent those impacted by kink formation. While the relative magnitude of RMSF varies, local extrema occur at consistent residues independent of simulation or protein for a given conformation. Beyond the kink in TM1, loops between helices in the transmembrane domain correlate with local maxima in RMSF. Note that the C-terminal domain tends to be more dynamic. (C) Per-residue RMSD measures the root mean square deviation of an individual residue from its initial position, averaged over the trajectory. Long form kink formation is clearly reflected by the extended region of elevated RMSD shaded cyan. Each result was gathered with the appropriate GROMACS command, using C_*α*_ atoms for all-atom simulations, and backbone beads for CG simulations. (D) Images of initial (partially transparent) and final frames for each single protein simulation, with kink location colored cyan. In agreement with (A)-(C), the short form maintains its structure, while a kink develops during all-atom and CG simulations of the long form. Lastly, as seen in the initial frame, kink development in the long form already appears to have started during the equilibration steps before production.

**Table S1.** Summary of molecular dynamics simulations performed in this study. The M protein(s) embedded is(are) specified in the second column, with total simulation time in the third, window for analysis in the fourth, and simulation purpose in the last. Restrained simulations are performed using the post-equilibrium system from the free counterpart.

| Category | System | Duration | Analysis window | Primary objective |
| --- | --- | --- | --- | --- |
| All-atom | Short form | 750 ns | 250–750 ns | Membrane deformation sign |
|  | Long form | 750 ns | 250–750 ns | Membrane deformation sign |
| Single protein<br>(coarse-grained) | Short form | 30 μs | 2–30 μs | Curvature field quantification |
| | Long form | 30 μs | 2–30 μs | Curvature field quantification |
| | Short form (restrained) | 10 μs | 2–10 μs | Curvature induction confirmation |
| | Long form (restrained) | 10 μs | 2–10 μs | Curvature induction confirmation |
| Protein pairs<br>(coarse-grained) | Short–short | 30 μs | 2–30 μs | Membrane-mediated interactions |
| | Long–short | 30 μs | 2–30 μs | Membrane-mediated interactions |
| | Long–long | 30 μs | 2–30 μs | Membrane-mediated interactions |

## Supplementary Text 2: Molecular dynamics minimization and equilibration

All-atom minimization and equilibration steps provided by CHARMM-GUI (59–65) began with initial energy minimization and then transitioned to two equilibration steps in the NVT ensemble using Berendsen temperature coupling (93) at 303.15 K. Following these were three further equilibration steps in the NPT ensemble, using both Berendsen temperature and Berendsen semi-isotropic pressure coupling. During minimization and equilibration, position restraints for lipids and protein atoms were gradually lessened before vanishing after the last equilibration step.

Alternatively, for our CG simulations, after an initial minimization step are three equilibration ones in the NPT ensemble, both using a Berendsen thermostat at 303.15 K and a semi-isotropic Berendsen barostat. The last two equilibration steps were also in the NPT ensemble with a v-rescale thermostat (77) at 303.15 K and a semi-isotropically applied c-rescale barostat (78). Restraints on lipids were used to prevent z-axis motion, while protein restraints were applied uniformly, with both gradually decreasing to zero after the end of equilibration.

## Supplementary Text 3: Principal curvature calculation

From the height polynomial generated using radial basis interpolation on average membrane midplane height grids, we use the shape operator (S) in the Monge gauge. With this parameterization (88), we have

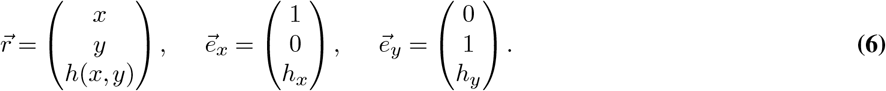

Where 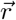 is the position vector, and 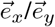 are the local tangent vectors. Note that *h*_*x,y*_ = ∂_*x,y*_*h*. As such, S is (87)

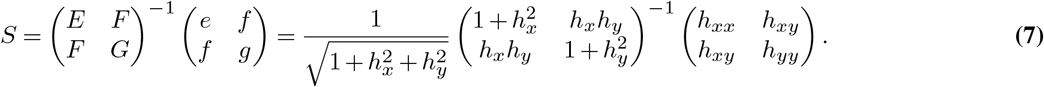

The eigenvalues of S correspond to our maximum (*k*_1_) and minimum (*k*_2_) curvatures, while the eigenvectors represent curvature directions (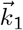 and 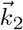),

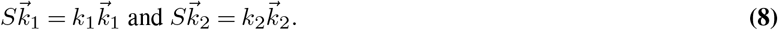

After finding 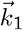 and 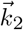 using *S* in the tangent plane, we then convert them to the original coordinate system for our final principal directions (Eq. 9),

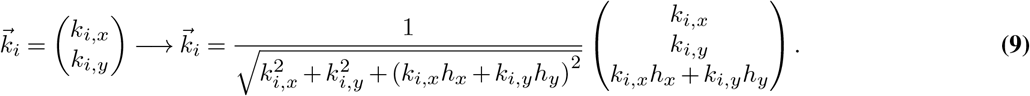

## Supplementary Text 4: Induced curvature averaging

Each protein’s induced maximum, minimum, total, or Gaussian curvature is defined as the average quantity adjacent to the protein’s average center of mass. The corresponding quantity of interest is taken at the four original grid points closest to the protein’s center to find this average, where each point is equidistant due to grid construction. For principal curvature directions, a circular average of 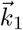 or 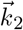 from these four points is used instead. This averaging convention is the simplest way to account for the finite protein cross-sectional area while also neglecting curvatures from interpolated points. However, while it is correct for the two intrinsic quantities (total and Gaussian curvature), averaging does not completely account for the possibility of each grid point having different directions of principal curvature in calculating induced maximum and minimum curvatures.

Full error (*σ*) estimation for these average quantities includes the standard deviation of the four points (*σ*_close_) and the intrinsic error of each involved point 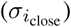,

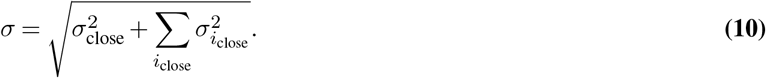

For all-atom systems, smoothing is the dominant component of intrinsic error. The impact of smoothing on each point is determined by re-generating the height and curvature for nine other Gaussian standard deviations ranging from 0.5 to 5 times the bin size. Standard deviation of these nine replicates at a point is the point’s intrinsic error.

Alternatively, intrinsic error for CG simulations is a combination of interpolation error and membrane midplane height standard error. As such, this quantity is calculated using Monte Carlo sampling, where the height of each point is defined by a normal distribution with mean equivalent to the original height, and standard deviation corresponding to the midplane standard error. Midplane standard error is found at every original grid point by determining - standard deviation of 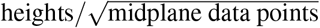 - for said point (this value is taken to be greater than or equal to the trajectory precision). Polynomials are generated according to 1000 different height combinations in CG systems, where the standard deviation of resulting values at a point is the point’s intrinsic error.

Intrinsic error is negligible for CG and substantial for all-atom systems. When curvature directions are involved, averages and standard deviation are converted to their circular counterparts. In comparing curvature magnitudes, short form error is larger than that of the long form. This stems from the more refined ridge-like shape of the short form’s deformation surrounding its center of mass, leading to an increased curvature variation across the protein’s cross-section.

